# Cortical α-synuclein pathology induces neuronal hypoactivity and circuit changes but not functional impairments in a model of early Lewy body dementia

**DOI:** 10.1101/2025.07.31.668024

**Authors:** Aye Theint Theint, Noe Cazares, Shujing Zhang, Ninglong Zhang, Rochelle Mosley, Kazuhisa Ishida, Makato Higuchi, Chao Peng, William Zeiger

## Abstract

Posterior cortical impairments in visuospatial/perceptual function are common non-motor symptoms of the Lewy Body Dementias (LBD) and are relatively specific for LBDs compared to other dementias. Across populations, cognitive impairments correlate with the presence of α-synuclein (α-syn) pathology in limbic and neocortical brain regions. However, the specific role that α-syn pathology plays in driving cortical circuit dysfunction and cognitive impairment remains controversial. We hypothesized that inducing α-syn pathology in primary visual cortex (V1) in mice would impair neuronal activity and encoding of visual information, leading to visuoperceptual impairments. To test this, we injected α-syn pre-formed fibrils (PFF) into V1, causing the formation of sparse Lewy-like pathology. Using longitudinal in vivo two-photon (2P) calcium imaging over 6 months, we recorded visually evoked activity of pyramidal cells in layer 2/3 (L2/3) and quantified α-syn pathology using C05-05, a fluorescent ligand that binds aggregated α-syn. Measuring population activity, we found a greater percentage of neurons in PFF-injected mice were responsive to visual stimuli with lower direction selectivity compared to controls at 4-5 months post-injection (MPI). Neurons with somatic Lewy-like inclusions showed reduced activity compared to neighboring neurons without inclusions. Conversely, neurons without somatic inclusions showed increased activity, positively correlated with the local burden of α-syn pathology. Using a coherent motion discrimination task, we found no impairments in visuoperceptual ability in PFF-injected mice. Our results demonstrate that α-syn pathology leads to reductions in neuronal activity in cells with somatic inclusions and reciprocal changes in local population activity but does not impair visuoperceptual function. Reflecting the early stages of neocortical α-syn pathology, our model provides a framework for future studies to better understand the heterogeneity of cognitive symptoms and α-syn pathology across patients.

## Introduction

The Lewy Body Dementias, including Parkinson’s disease dementia (PDD) and DLB, are common neurodegenerative disorders with both motor and non-motor symptoms^1–3^. Cognitive impairment is a defining feature of DLB and is also common in PD. For example, 50–80% of patients with PD develop cognitive impairment within 10–20 years of diagnosis^4–7^. The most commonly affected cognitive domains include attention, executive function, and visuospatial/perceptual processing^2,6,8–10^. These impairments likely result from dysfunction within specific neural circuits^9,11^. Executive dysfunction is often linked to frontostriatal circuits^12,13^ and is partly dependent on dopaminergic signaling^14–16^. These deficits are nearly ubiquitous amongst PD patients and do not strongly correlate with, or predict progression to, dementia^9^. In contrast, visuospatial and perceptual deficits are associated with dysfunction in posterior cortical regions such as the parietal, temporal, and occipital lobes, where patients show selective hypometabolism and cortical atrophy^17–19^. These posterior cortical changes are closely linked to cognitive decline and are predictive of dementia in PD^9,17,19,20^. Therefore, dysfunction in posterior cortical circuits may play a critical role in visuospatial and perceptual deficits and the development of dementia in PD and DLB. However, the underlying mechanisms remain poorly understood.

PD and DLB are pathologically defined by the progressive accumulation of α-syn into insoluble intraneuronal inclusions called Lewy bodies and Lewy neurites^21–23^. Neuropathological studies have demonstrated a correlation between the presence of α-syn pathology, disease severity, and specific symptoms. For instance, α-syn pathology is often found in brainstem nuclei such as the dorsal motor nucleus of the vagus and the substantia nigra early in the disease course^24^. Pathology in these regions correlates with prodromal symptoms such as REM sleep behavior disorder and early motor impairments. As disease progresses and becomes more severe, α-syn pathology is found more frequently in limbic and neocortical regions where it correlates with cognitive impairment and dementia^25–27^. Diffuse neocortical α-syn pathology is also the hallmark for the pathological diagnosis of DLB^2,28^. While these correlations are compelling, at the level of individuals, patients with PD have been reported with substantial neocortical α-syn burdens and minimal cognitive impairments, or minimal neocortical α-syn and substantial cognitive impairment^25,26,29–31^. Thus, it remains debated whether, and to what extent, α-syn pathology drives cortical circuit dysfunction and symptoms of cognitive impairment.

In vitro, overexpression or aggregation of α-syn has been reported to disrupt synaptic vesicle recycling and to impair excitatory neurotransmission^32–34^. To date, only a limited number of studies have been conducted to directly assess the impact of α-syn pathology on cortical neuron or circuit activity in vivo. These studies have demonstrated selective degeneration of neurons harboring inclusions^35^, reduction in dendritic spine density^36^, alterations in calcium transients^37^, and increased whisking- and locomotion-evoked activity^38,39^. However, none of these studies examined circuit changes in cortical areas relevant for cognitive impairment in PD, no longitudinal data exists on how posterior cortical circuits are affected throughout disease progression, and it has not been possible to test the specific relationship between altered neuronal activity and the presence or absence of α-syn pathology in individual neurons. Resolving the specific role of α-syn pathology in driving cognitive impairments is crucial as clinical trials targeted at reducing α-syn as a disease modifying strategy for cognitive impairment in PD (ClinicalTrials.gov ID: NCT07055087) are currently in process.

Given the limited data currently available, we hypothesized that cortical α-syn pathology directly impairs neuronal activity and circuit function, and that these effects would evolve throughout disease progression. To test this, we utilized the PFF model of PD associated pathology. We injected α-syn PFFs in V1 given the importance of posterior cortical circuit dysfunction in the development of visuoperceptual deficits, cognitive impairment, and dementia. We then performed longitudinal 2P in vivo calcium imaging of neuronal activity for up to 6 MPI. We also utilized C05-05^40^, a green fluorescent ligand that specifically binds aggregated α-syn, to visualize α-syn pathology in vivo with single neuron resolution. Injection of PFFs in V1 seeds sparse α-syn pathology that evolves over time in V1 and spreads to other brain regions. Compared to controls, mice injected with PFFs show small but significant changes in spontaneous and visually evoked activity at the population level, peaking at 4-5 MPI. At the level of single cells, neurons containing somatic α-syn aggregates are less active than surrounding neurons, and greater intracellular pathology burden correlates with lower activity. In contrast, neighboring neurons without aggregates tend to be more active, with greater local α-syn burden correlated with increased activity. Using a translationally relevant visuoperceptual motion discrimination assay^41^, we found that visual function remains preserved in PFF-injected mice up to 6 MPI. Our results demonstrate, for the first time in vivo, hypoactivity of neurons with α-syn aggregates. Our data also identify changes in local population activity that could be compensatory helping to preserve visuoperceptual function. These findings support a direct role for α-syn in cortical circuit changes in a model of early neocortical PD-associated pathology, but suggest that other mechanisms are likely necessary to drive functional cognitive decline in LBD.

## Results

### A novel model of sparse posterior cortical α-syn pathology

To understand how α-syn pathology affects cortical circuit function, we developed a model in which we could reliably seed α-syn pathology in the cortex. We chose to focus on V1 given visuoperceptual impairments are common in patients with PD and DLB^10^, and because posterior cortical circuit dysfunction is linked to the development of dementia^9,20^. We injected mouse PFFs directly into V1 to concentrate pathology in visual cortical areas and to avoid potential confounds of dopaminergic neuron loss that can be seen with striatal injections^60^. After injection, we performed immunofluorescence staining for α-syn phosphorylated at S129 (pS129) to assess the distribution of α-syn pathology at 1, 3, and 6 MPI. At 3 MPI, we observed pS129^+^ α-syn pathology throughout V1 (**Fig. 1a**). Pathology was enriched in specific cortical layers, especially layer 5 (L5) and layer 6a (L6a), and spread to the contralateral V1 and other cortical and sub-cortical regions (**Fig. 1b-d**). A similar laminar distribution was observed when we included higher visual areas in addition to V1 (**Fig. S1a**). Over time, we found pathology in V1 increased in density, peaking between 3 and 6 MPI (**Fig. 1d, Fig. S1a**), similar to the time course observed for striatal PFF injections^61^. Even at 3 MPI, the density of pathology was relatively sparse, in agreement with neuropathology data from visual cortical areas in patients with PD and DLB^62–64^. Other brain regions with notable pS129 staining included the ipsilateral dorsomedial striatum, hippocampus, and secondary motor (M2)/prefrontal areas (**Fig. 1d, Fig. S1b-c**). The time course for development of pathology was generally similar in these areas, peaking at 3 to 6 MPI. Interestingly, we did not observe any pS129 staining in the dorsolateral geniculate nucleus (dLGN), which provides strong afferent inputs to V1 (**Fig. 1d**). Immunofluorescence analysis showed that pS129^+^ α-syn aggregates were predominantly localized within CaMKII^+^ excitatory neurons at 1, 3, and 6 MPI, with little to no colocalization observed in Iba1^+^ microglia (**Fig. S2)**. Furthermore, while robust pS129 pathology was detected only in PFF-injected mice, the distribution and morphology of Iba1^+^ microglia and GFAP^+^ astrocytes were qualitatively comparable among PFF-, α-syn monomer-, and saline-injected groups, indicating no overt neuroinflammatory responses associated with PFF injections (**Fig. S3**, **Fig. S4**). Thus, injection of PFFs into V1 seeds α-syn pathology that develops over time and spreads to anatomically connected brain regions.

**Figure 1:**
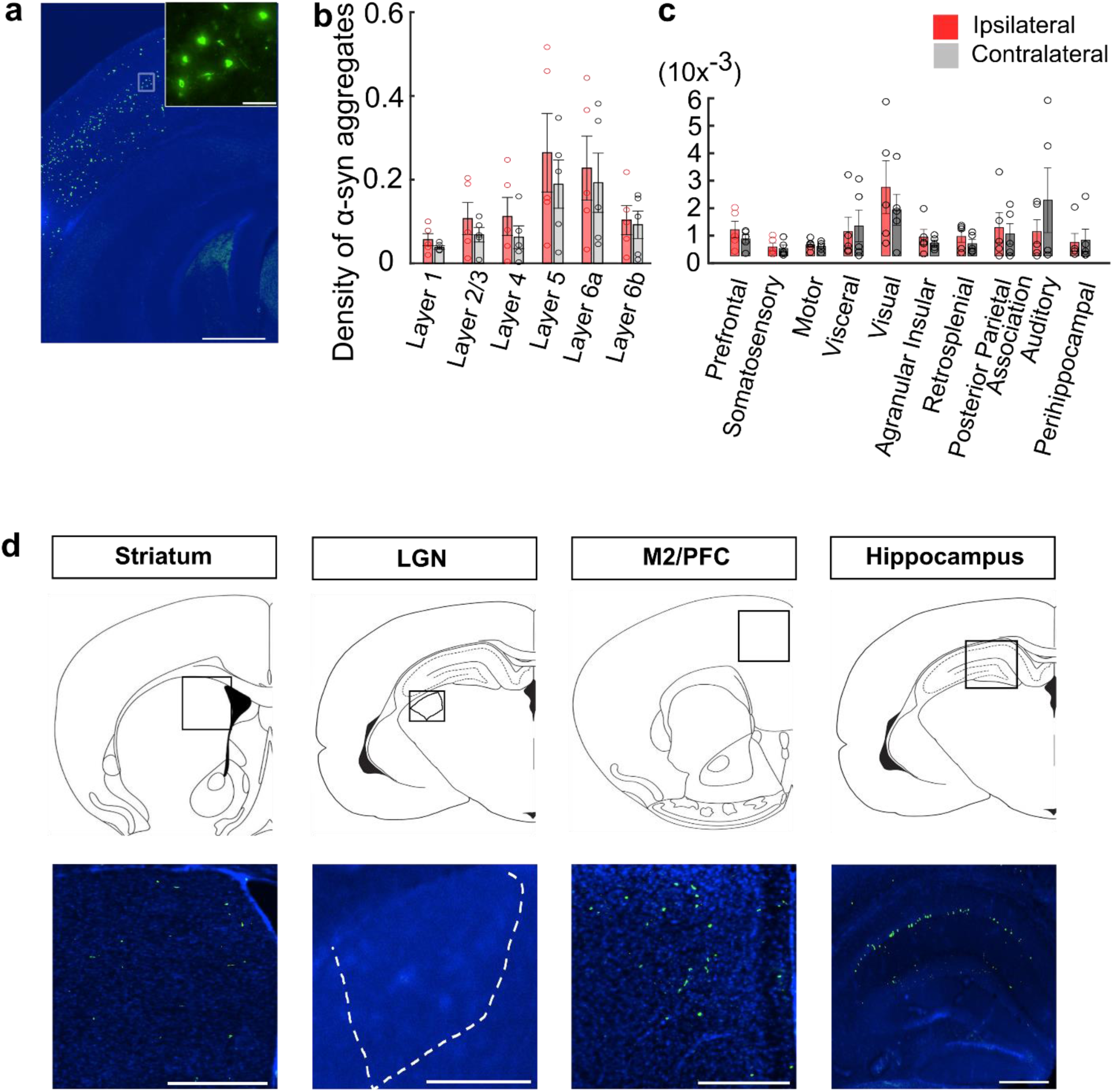
Injection of PFFs in V1 seeds pS129^+^ α-syn pathology in mice. **a:** Representative image showing pS129^+^ α-syn aggregates (green) together with DAPI counterstain (blue) in V1 from a coronal brain section from a PFF-injected mouse at 3 MPI. Scale bar: 500 µm (inset from boxed area, scale bar: 50 µm). **b-c**: Density of pS129^+^ α-syn aggregates in each layer of ipsilateral (red) and contralateral (gray) V1 (*b*) and other cortical regions (*c*) at 3 MPI. Bars show mean ± s.e.m., with data points for individual mice (n=5 mice) overlaid. **d:** Schematic cartoons of coronal brain sections with boxes indicating the regions in the corresponding images below (Top row). Representative regions with pS129^+^ α-syn aggregates in coronal brain sections from mice injected with PFFs in V1 at 3 MPI. pS129^+^ α-syn aggregates (green) are visible with DAPI counterstain (blue) (Bottom row). Scale bar: 200 µm. LGN = lateral geniculate nucleus, shown in the outlined region; M2 = secondary motor cortex; PFC = prefrontal cortex.

### Progression of α-syn pathology is associated with changes in population activity of L2/3 neurons in V1

To determine how the development of α-syn pathology affects posterior cortical circuit function, we injected PFFs into the left V1 of 7–12-week-old mice stably expressing the red-shifted genetically encoded calcium indicator jRGECO1a in excitatory cortical neurons (Thy1-jRGECO1a mice). Control mice were injected with the same volume of PFF vehicle (phosphate buffered saline). We also implanted a glass cranial window over V1 to facilitate in vivo imaging. Next, we localized V1 using intrinsic optical signal imaging (IOSI, **Fig. S5a**) and then performed 2P in vivo imaging of calcium signals evoked by drifting sinusoidal gratings in L2/3 of awake mice at monthly intervals up to 6 MPI (**Fig. 2a-c**). We did not observe filled cells typical of calcium indicator toxicity at any time up to 6 MPI, consistent with prior reports showing long-term stability of jRGECO1a expression in these mice^65^ (**Fig. S6**). Drifting grating stimuli spanned 8 directions (from 0 to 315°, at 45° intervals) and were presented in pseudorandom order to the right eye contralateral to the PFF injection site a total of 4 times for each direction. For all active neurons, we extracted calcium fluorescence traces and calculated modified Z-scores to facilitate comparisons across neurons with differing baseline fluorescence levels (**Fig. S6b-c**). We then quantified the area-under-the-curve (AUC) of evoked responses to individual stimuli, the mean evoked response for each grating direction, direction selectivity (DSI), and orientation selectivity (OSI) (**Fig. 2d-e**). Neurons were classified as visually responsive by determining if their activity trace was significantly correlated to a trace of the displayed visual stimuli (see materials and methods). At 1 MPI we found that 38% of neurons were visually responsive in both control and PFF-injected mice (38.6 ± 2.1% vs. 38.7 ± 2.1%, respectively; **Fig. 2f**), similar to the percentage of visually responsive neurons observed in other studies using jRGECO1a^65,66^. To assess for changes within and between groups over time, and to account for clustering of observations from hundreds of neurons within individual mice, we analyzed our data using linear mixed effects models^67^. We found that over time there was a small but significant decrease in the percentage of visually responsive neurons in both groups of mice (**Fig. 2f**). Interestingly, there was also a significant group-by-timepoint interaction, with PFF-injected mice having a greater percentage of visually responsive neurons compared to saline-injected controls at 2, 4, and 5 MPI (**Fig. 2f**). The difference was greatest at 4 MPI, with PFF-injected mice showing a 6.55% increase in visually responsive neurons compared to controls (38.9% vs. 32.4%, respectively).

**Figure 2:**
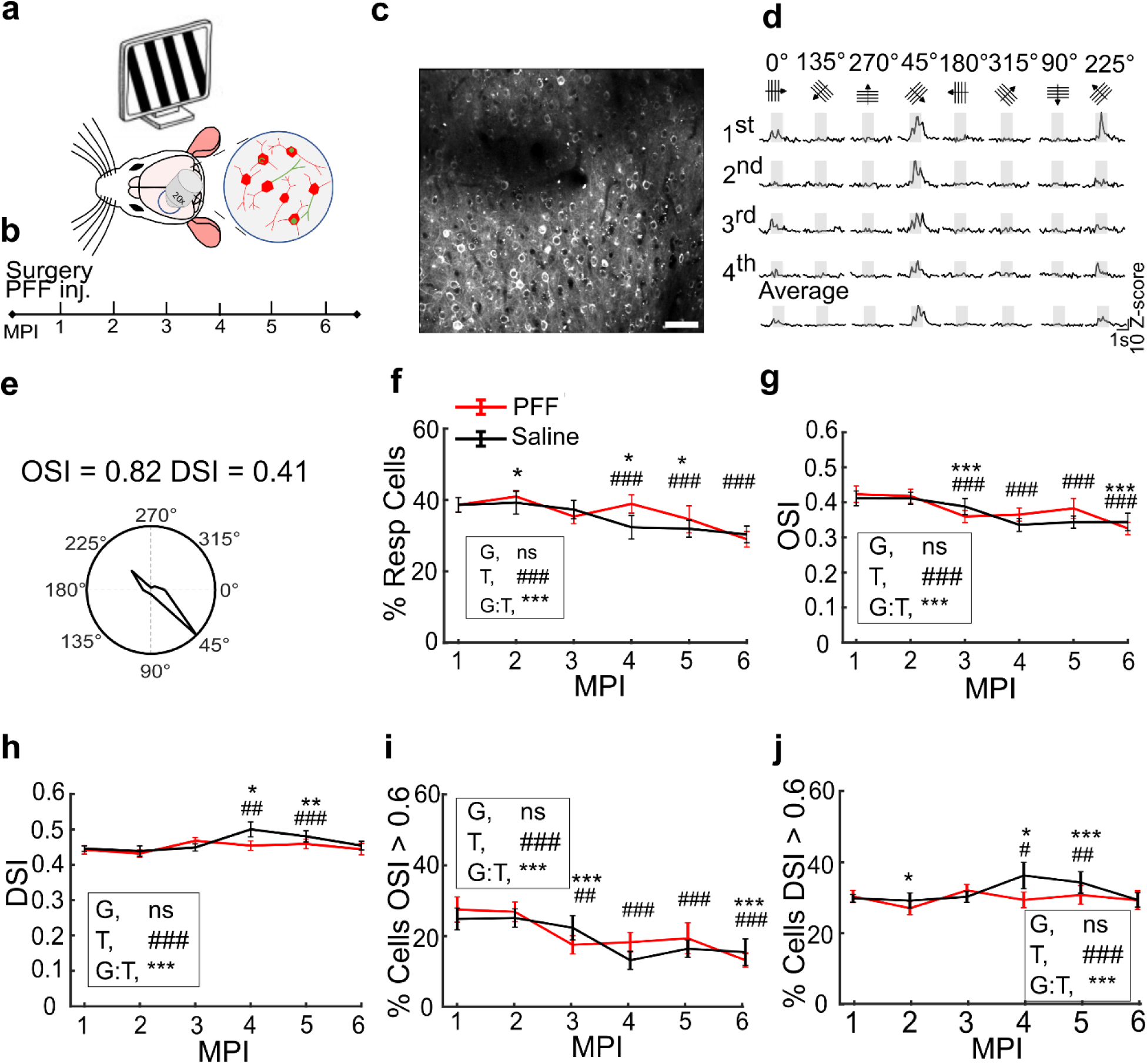
α-syn pathology causes subtle changes in visually evoked activity of L2/3 neurons over time. **a**: Schematic of 2P imaging of L2/3 neurons labeled with jRGECO1a (red) and C05-05 (green). **b**: Experimental timeline. **c**: Average projection image of L2/3 neurons at 1 MPI. Scale bar = 50 μm. **d**: Example of stimulus evoked calcium transients from one neuron. Orientation and direction of stimuli are indicated at the top. Gray shading indicates periods of visual stimuli (1 s long, 3 s interstimulus interval). Four individual trials with the trial averaged responses are shown for each stimulus. **e**: Polar plot summarizing normalized stimulus evoked responses from the same neuron from *d*. **f**: Percentage of visually responsive L2/3 neurons in V1 for control (gray) and PFF-injected (red) mice. GLME binomial model, ANOVA for fixed effects of group (G, *p* = 0.896), timepoint (T, ###, *p* = 7.11 × 10^−15^), and group-by-timepoint interaction (G:T, ***, *p* = 3.47 × 10^−6^). **g**: Mean orientation selective index (OSI) of L2/3 neurons in V1 for control and PFF-injected mice. LME model, ANOVA for fixed effects of group (G, $ *p* = 0.385), timepoint (T, ###, *p* = 2.17 × 10^−25^), and group-by-timepoint interaction (G:T, ***, *p* = 2.86 × 10^−8^). **h**: Mean direction selective index (DSI) of L2/3 neurons in V1 for control and PFF-injected mice. LME model, ANOVA for fixed effects of group (G, *p* = 0.637), timepoint (T, ###, *p* = 4.04 × 10^−6^), and group-by-timepoint interaction (G:T, **, *p* = 2.58 × 10^−4^). **i**: Percentage of responsive cells with OSI > 0.6 for control and PFF-injected mice. GLME binomial model, ANOVA for fixed effects of group (G, *p* = 0.324), timepoint (T, ###, *p* = 5.32 × 10^−12^), and group-by-timepoint interaction (G:T, ***, *p* = 5.87 × 10^−7^). **j**: Percentage of responsive cells with DSI > 0.6 for control and PFF-injected mice. GLME binomial model, ANOVA for fixed effects of group (G, *p* = 0.406), timepoint (T, ###, *p* = 3.10 × 10^−3^), and group-by-timepoint interaction (G:T, ***, *p* = 6.42 × 10^−4^).

The mean orientation selectivity and direction selectivity of neurons in control mice showed some small, variable changes over time. There were also significant group-by-timepoint interactions, with PFF-injected mice showing lower orientation selectivity at 3 and 6 MPI and lower direction selectivity at 4 and 5 MPI (**Fig. 2g-h**). Overall tuning width trended up slightly over time in both groups but was largely similar between control and PFF-injected mice (**Fig. S5b**). We also quantified the percentage of highly orientation and direction selective cells (OSI/DSI > 0.6) in control and PFF-injected mice over time. The proportion of these highly selective cells changed in a similar fashion to the changes observed in mean OSI and DSI, with fewer cells with OSI>0.6 at 3 and 6 MPI, and fewer cells with DSI>0.6 at 2, 4 and 5 MPI (**Fig. 2i-j, Fig. S5c-n**). Together, these results demonstrate that the development of cortical α-syn pathology leads to time-dependent changes in visually evoked activity of L2/3 neurons at the population level, with PFF-injected mice showing more visually responsive neurons with lower orientation and direction selectivity.

We next investigated “spontaneous” neuronal activity in mice during quiet wakefulness in the absence of visual stimuli. We recorded activity for 5 minutes and calculated the AUC of the entire calcium trace as a measure of total spontaneous activity, as well as the frequency and amplitude of individual calcium transients (**Fig. 3a**). The AUC of both groups was similar at 1 MPI (212.9 ± 5.7 Z-score*sec in control vs. 220.8 ± 5.9 Z-score*sec in PFF-injected mice) and there were small fluctuations in the control group over time (**Fig. 3b**). There was a significant group-by-time interaction with PFF-injected mice showing trends for lower AUC, though this was only significantly different from controls at 3 MPI (**Fig. 3b**). Calcium transient frequency fluctuated slightly over time in control animals, but there were small, significant group-by-time differences in PFF-injected mice, with lower frequencies at 2 and 3 MPI and higher frequencies at 4 and 5 MPI compared to controls (**Fig. 3c**). The mean amplitudes of calcium transients showed very small fluctuations over time in the control group and significant group-by-time interactions from 3-6 MPI, with the PFF-injected mice showing lower amplitudes, though the magnitude of this change was very small (**Fig. 3d**).

**Figure 3:**
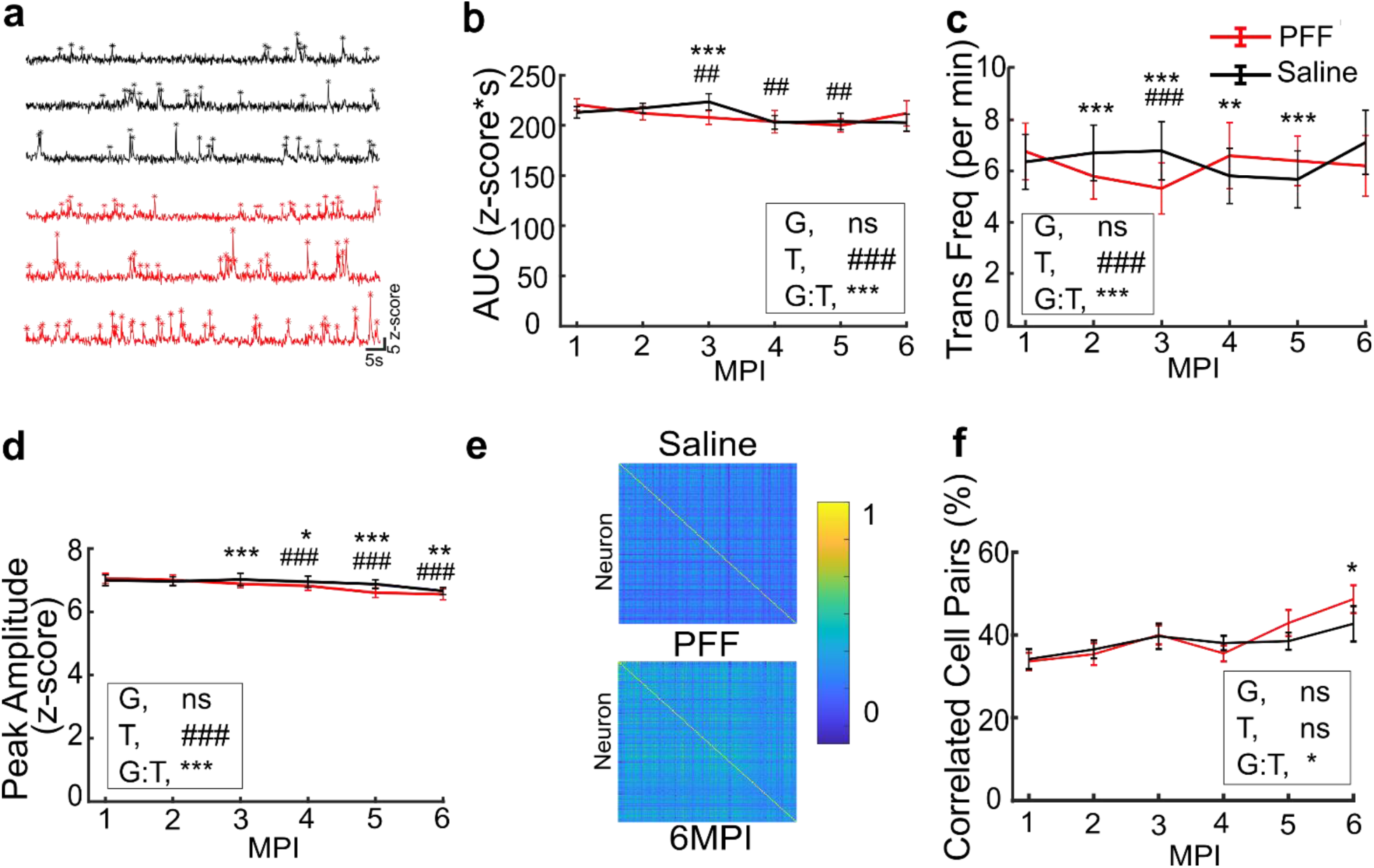
α-syn pathology causes subtle changes in spontaneous activity of L2/3 neurons in V1 over time. **a:** Representative z-scored jRGECO1a fluorescence signal traces from control (black) and PFF-injected (red) mice at 3 MPI. Asterisks denote detected calcium transient peaks used for quantification in *c* and *d*. **b:** Mean AUC from the entire spontaneous activity trace from neurons in PFF-injected or control mice. LME model, ANOVA for fixed effects of group (G, *p* = 0.761), timepoint (T, ###, *p* = 5.08 × 10^−10^), and group-by-timepoint interaction (G:T, ***, *p* = 7.09 × 10^−9^). Significance for individual coefficients for timepoint (T) or group-by-timepoint interaction (G:T), corrected using B & H method, are indicated over corresponding data points in *b-d,f* (# and *, *p* < 0.05; ## and **, *p* < 0.01; ### and ***, *p* < 0.001). Individual timepoints are means ± s.e.m for data in *b-d,f.* **c:** Mean frequency of calcium transients (per minute) from PFF-injected or control mice. LME model, ANOVA for fixed effects of group (G, *p* = 0.954), timepoint (T, ###, *p* = 1.21 × 10^−4^), and group-by-timepoint interaction (G:T, ***, *p* = 2.78× 10^−51^). **d:** Mean peak amplitude of calcium transients from PFF-injected or control mice. LME model, ANOVA for fixed effects of group (G, *p* = 0.929), timepoint (T, ###, *p* = 6.91 × 10^−27^), and group-by-timepoint interaction (G:T, ***, *p* = 5.75× 10^−15^). **e:** Correlograms of pairwise correlations of calcium transients from PFF-injected or control mice at 6 MPI. The scale denotes the Pearson correlation coefficient. **f:** Mean percentage of correlated cell pairs from PFF-injected or control mice. LME model, ANOVA for fixed effects of group (G, *p* = 0.595), timepoint (T, *p* = 0.117), and group-by-timepoint interaction (G:T, *, *p* = 0.0489).

In addition to the amount of total activity, the timing of coordinated neuronal activity is essential for sensory coding^68^. Changes in synchronized activity provide information about functional connectivity between neurons^68,69^, and alterations in neuronal synchrony have been observed in mouse models of other neurodegenerative diseases, for example, models of Aβ deposition^70,71^. Therefore, we investigated synchrony of neuronal activity in PFF-injected mice. First, we quantified the percentage of pairs of active neurons with significantly correlated activity within each imaging field-of-view (FOV) during spontaneous activity. The percentage of correlated cell pairs was relatively stable in control mice, without significant changes over time (**Fig. 3e-f**). There was a significant group-by-time interaction, with PFF-injected mice showing an increase in the number of correlated cell pairs as pathology progressed, with a statistically significant increase observed at 6 MPI (**Fig. 3e-f**). A recent study reported that locomotion-induced synchrony of neurons in the primary motor cortex (M1) is increased in mice injected with PFFs in M1 or the dorsolateral striatum^39^. To assess for potential region-specific effects of PFFs, we analyzed neuronal synchrony using similar methods, quantifying pairwise synchronicity and ensemble stability during visually evoked neuronal activity. Control animals showed a slight increase in pairwise synchronicity over time (**Fig. S7a-b**). There were significant group-by-time differences in PFF-injected mice, but these were inconsistent over time and of very small magnitude, unlikely to biologically meaningful (**Fig. S7b-c**). Ensemble stability was slightly lower in PFF-injected mice starting at 1 MPI with no change over time in either group (**Fig. S7c**).

### Effects of somatic α-syn inclusions on neuronal activity in V1

Given that cortical α-syn pathology is sparse, and we observed highly heterogenous neuronal responses among individual cells, we hypothesized that the effects of α-syn might be different in cells with or without somatic α-syn inclusions. To assess for potential cell-autonomous impacts of α-syn pathology, we used C05-05, a fluorescent ligand that binds specifically to aggregated α-syn^40^, to label and image α-syn inclusions in vivo at 4-5 MPI, at the peak of α-syn pathology density. To validate the specificity of C05-05 labeling, we performed co-immunostaining with an anti-pS129 antibody. Qualitatively we found strong overlap between C05-05 signals and pS129 staining (**Fig. S8**), consistent with our previous results^40^. A quantitative object-based analysis further demonstrated substantial colocalization between the two signals, with 58.4% of all pS129^+^ objects overlapping at least partially with C05-05^+^ signal. Among these colocalized objects, the mean overlap fraction, defined as the proportion of the pS129^+^ object area overlapping with C05-05 signal, was 68.5%. In contrast, only 10.2% of C05-05^+^ objects were false positives, showing no overlap with pS129^+^ objects, and these tended to be small weakly fluorescent signals (**Fig. S8**). Together, these results validate the specificity of C05-05 labeling, particularly for larger α-syn inclusions.

In vivo, we found the emission spectra of C05-05 (**Fig. S9a**) led to clearly separable signals for C05-05 and jRGECO1a, with no significant bleed through between channels (**Fig. S9b**). We manually scored and identified neurons containing clear, somatic C05-05 positive (C05-05^+^) inclusions (**Fig. 4a**). We also selected multiple C05-05 negative (C05-05^-^) neurons within 200 μm of each C05-05^+^ neuron in the same FOV. C05-05^+^ neurons exhibited markedly lower activity compared to C05-05⁻ neurons across a range of activity parameters, including lower mean z-scored activity (0.746 ± 0.825 vs. 1.10 ± 0.84 z-score, **Fig. 4b**) and lower AUC (116.5 ± 11.73 vs. 166.1 ± 10.1 z-score*sec, **Fig. 4c**) across the entire visually evoked imaging session, lower calcium transient frequency (10.5 ± 9.6 vs. 15.5 ± 8.9 transients per min, **Fig. 4d**), and lower calcium transient amplitude (6.76 ± 2.15 vs. 7.53 ± 1.72 z-score, **Fig. 4e**). We then compared specific visually evoked activity of C05-05^+^ and C05-05^-^ neurons. The proportion of visually responsive neurons was similar between groups, with 32% (32 out of 100) of C05-05^+^ neurons and 28.7% (44 out of 153) of C05-05^-^ neurons visually responsive (**Fig. 4g**). The DSI of C05-05^+^ (0.432 ± 0.050) and C05-05^-^ (0.513±0.040) neurons was similar (**Fig. 4h**), but the OSI of C05-05^+^ neurons (0.204 ± 0.031) was slightly, but significantly lower than that of C05-05^-^ neurons (0.303 ± 0.033, **Fig. 4i**). There was no difference in the modulation index (MI) of C05-05^+^ (1.22 ± 0.20) and C05-05^-^ (1.01 ± 0.13) neurons at their preferred orientation (**Fig. 4j**). We found similar lower activity of C05-05^+^ neurons compared to C05-05⁻ neurons in spontaneous conditions across a range of activity parameters, including lower mean z-scored activity (0.886 ± 0.075 vs. 1.19 ± 0.07 z-score, **Fig. S10a**) and lower AUC (273.2 ± 20.3 vs. 356.8 ± 20.5 z-score*sec, **Fig. S10b**) across the entire visually evoked imaging session, lower calcium transient frequency (23.4 ± 1.1 vs. 26.4 ± 0.9 transients per min, **Fig. S10c**), and lower calcium transient amplitude (5.41 ± 0.11 vs. 5.83 ± 0.12 z-score, **Fig. S10d**).

**Figure 4:**
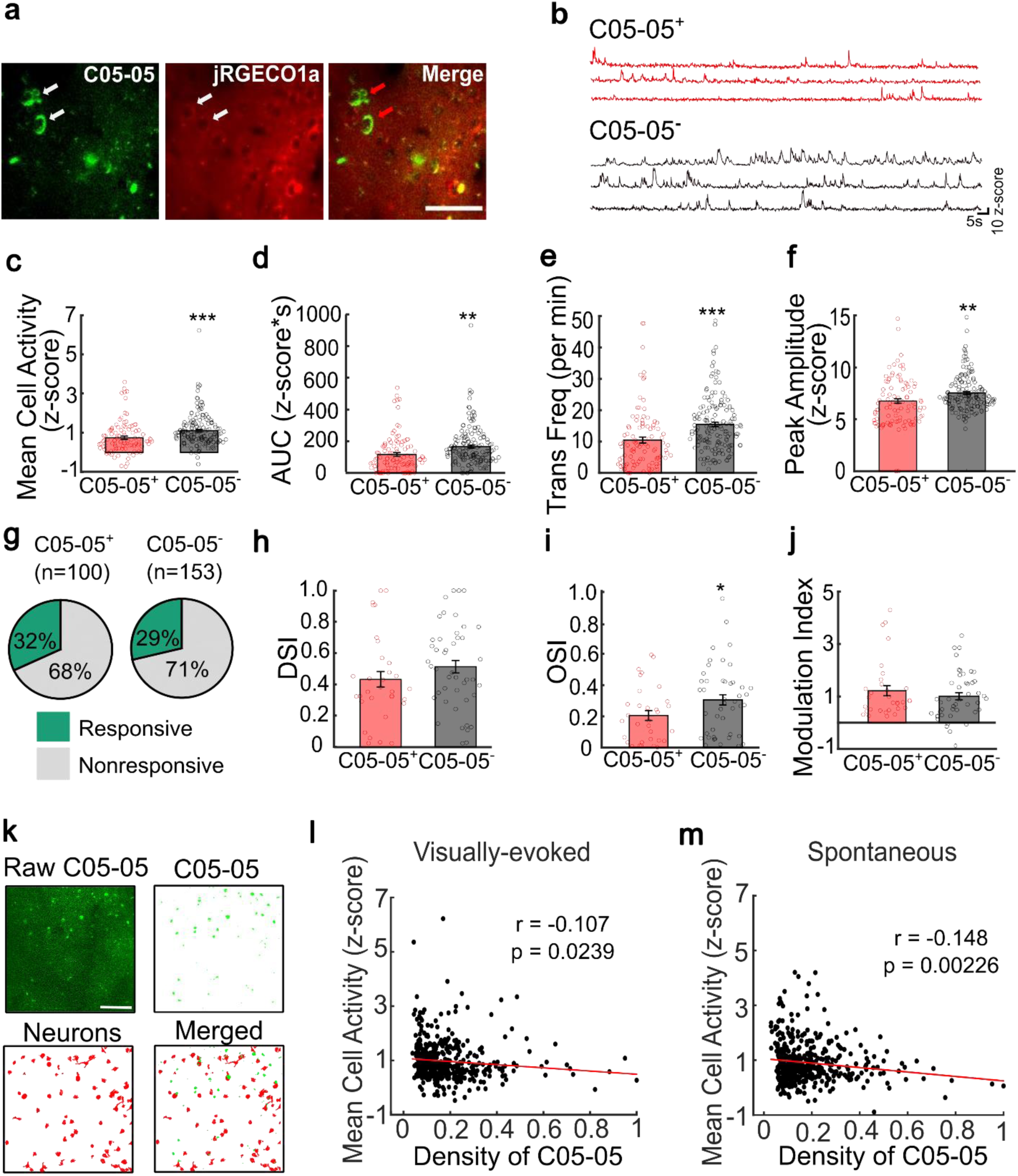
Intracellular α-syn inclusions are associated with hypoactivity of neurons in V1 of PFF-injected mice. **a:** 2P images of C05-05 (left), jRGECO1a (middle) and the composite image (right) in a PFF-injected mouse at 4 MPI. Arrowheads indicate jRGECO1a^+^ neurons with C05-05^+^ somatic α-syn inclusions. Scale bar = 50 μm. **b:** Representative z-scored jRGECO1a traces from C05-05^+^ (red) and C05-05^-^ (black) neurons. **c-f:** Mean z-scored fluorescence (*c*), AUC (*d*), transient frequency (*e*), and transient peak amplitude (*f*) in C05-05^+^ (red, n = 100 cells) and C05-05^-^ (gray, n = 153 cells) neurons from PFF-injected mice at 4 and 5 MPI. Two-sample, two-tailed *t* test, *p* = 9.60 × 10^−4^ (*c*), *p* = 0.0017 (*d*), *p* = 3.63 × 10^−5^ (*e*) and *p* = 0.002 (*f*). Dots shown for individual neurons. **g:** Proportion of visually responsive C05-05^+^ (left) and C05-05^-^ (right) neurons. **h:** Mean DSI of visually responsive C05-05^+^ (red, n = 32 cells) and C05-05^-^ (gray, n = 44 cells) neurons at 4 and 5 MPI. Two-sample, two-tailed *t* test, *p* = 0.207. **i:** Mean OSI of visually responsive C05-05^+^ (red, n = 32 cells) and C05-05^-^ (gray, n = 44 cells) neurons at 4 and 5 MPI. Two-sample, two-tailed *t* test, *p* = 0.0394. **j:** Modulation index (MI) to the preferred orientation of visually responsive C05-05^+^ (red, n = 32 cells) and C05-05^-^ (gray, n = 44 cells) neurons. Two-sample, two-tailed *t* test, *p* = 0.3626. **k:** Workflow for automated C05-05 scoring. C05-05 fluorescence (top left) was binarized (top right) and then merged with jRGECO1a segmentation maps (bottom left) to quantify C05-05 signals in individual neurons (bottom right). Scale bar = 100 μm. **l-m:** Correlation between mean z-scored calcium fluorescence of C05-05^+^/jRGECO1a^+^ neurons and somatic C05-05 density in visually evoked (*l*) and spontaneous (*m*) conditions. Pearson correlation coefficient (r = −0.107, *p* = 0.0239, n = 442 neurons in *h* and r = −0.148, *p* = 0.00226, n = 426 neurons in *m*).

To overcome potential biases in manual scoring of neurons for C05-05^+^ inclusions, we also implemented an automated method for C05-05 scoring. We binarized and segmented C05-05 fluorescence signals and overlaid them onto jRGECO1a^+^ fluorescence signals segmented by Suite2P (**Fig. 4k**). We then set a threshold of 10 pixels to differentiate cells with clear somatic C05-05^+^ inclusions apart from those with background signals. Plotting the mean activity of each cell compared to the density of C05-05 signal in the soma, we found a significant negative correlation between C05-05 burden and both visually evoked (**Fig. 4l**) and spontaneous (**Fig. 4m**) activity. To test for the possibility that C05-05 itself could alter jRGECO1a calcium signals and thereby confound measurements of neuronal activity, we recorded calcium activity in primary neuronal cultures expressing jRGECO1a before and after C05-05 exposure. Selecting cells with somatic C05-05^+^ inclusions for analysis, we found no significant changes in mean z-scored activity, AUC, calcium transient frequency or amplitude after C05-05 exposure (**Fig. S11**). This confirms that the reduced activity observed in C05-05^+^ neurons in vivo is unlikely to be an artifact of C05-05 binding to α-syn inclusions on jRGECO1a dynamics. Collectively, these findings demonstrate that α-syn pathology affects the activity of neurons within the visual cortex, inducing hypoactivity in neurons that correlates with the burden of intracellular somatic α-syn inclusions.

### Neurons without somatic α-syn inclusions show increased neuronal activity that correlates with the local burden of α-syn

Although we found an association between reduced activity and somatic α-syn inclusions, most neurons in L2/3 did not have somatic α-syn inclusions. Therefore, to determine how α-syn pathology affects circuit function in V1 more broadly, we next investigated how the presence of α-syn pathology within the local neuronal microenvironment influences the activity of surrounding neurons without somatic inclusions. Using the same approach described in Figure 4g, we quantified the total C05-05 signal in FOVs from mice 4-5 MPI and then correlated this with the mean activity of C05-05^-^ neurons in the same FOV (**Fig. 5a**). Interestingly, we found a positive correlation, with higher local burden of C05-05 signal associated with greater mean activity of neurons, in both visually evoked (**Fig. 5b**) and spontaneous (**Fig. 5c**) conditions. To determine if this relationship exists at the single neuron level, we identified C05-05^-^ neurons and quantified the total surrounding C05-05 signal within a radius of 200 pixels (166 µm). We then correlated mean activity of individual C05-05^-^neurons with the local C05-05 burden and again found a positive correlation, with greater visually evoked (**Fig. 5e**) and spontaneous (**Fig. 5f**) activity associated with more surrounding C05-05 signal. This relationship also held over smaller distances (within a radius of 100 pixels [83 µm], **Fig. S12**). Together, these findings suggest that as the local burden of α-syn pathology increases, the activity of neurons without somatic α-syn inclusions increases.

**Figure 5:**
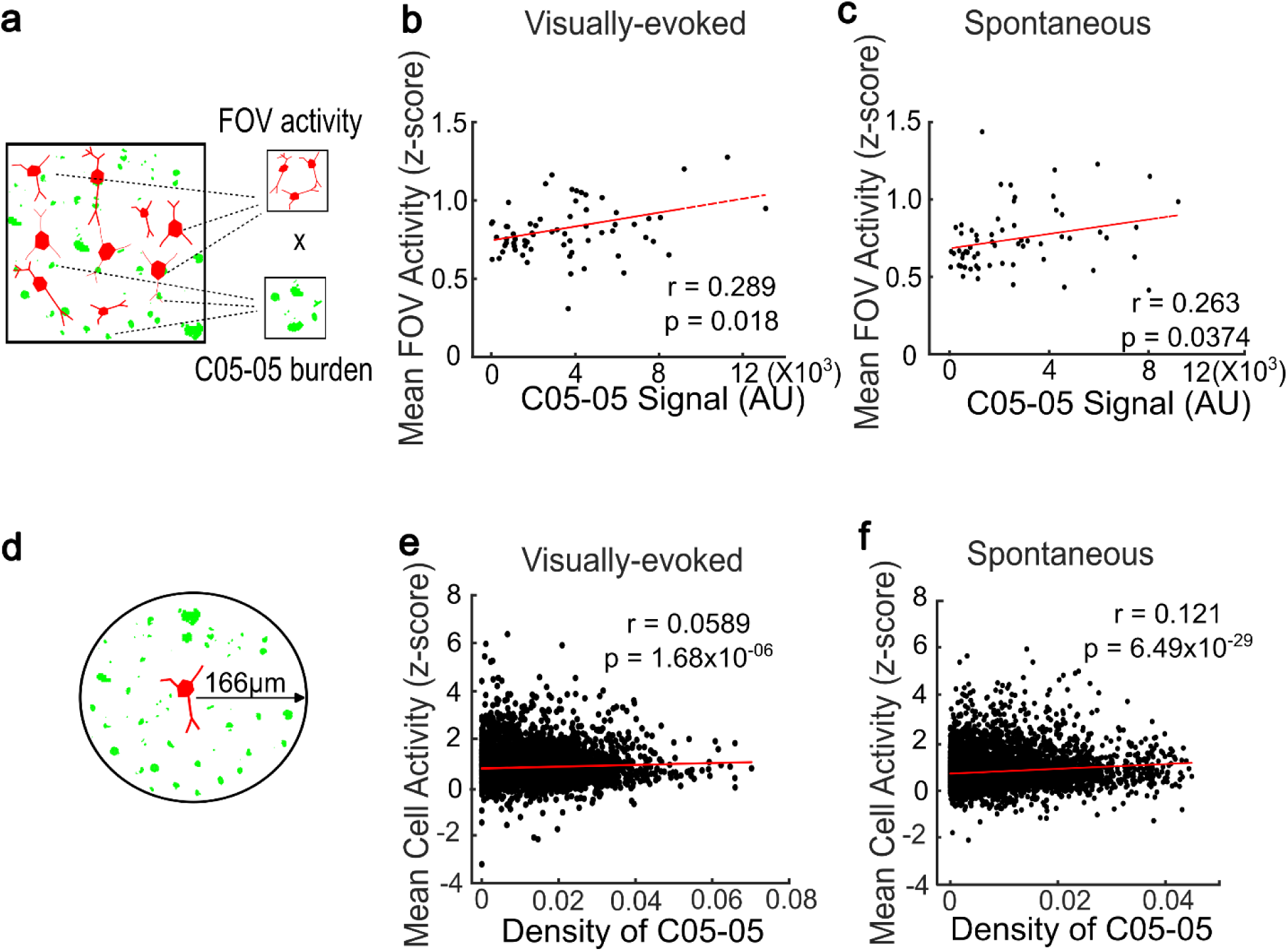
Local burden of α-syn pathology is associated with increased activity of inclusion-free neurons in V1. **a:** Schematic depicting correlation between mean population activity and C05-05 burden in an imaging FOV. **b-c:** Correlation between mean z-score of calcium fluorescence traces of C05-05^-^ neurons and total burden of C05-05 signal in a FOV from PFF-injected mice under visually evoked (*b*) and spontaneous (*c*) conditions at 4-5 MPI. Pearson correlation coefficient (r = 0.289, *p* = 0.018, n = 67 FOVs in *b* and r = 0.263, *p* = 0.0374, n = 63 FOVs in *c*). **d:** Schematic depicting quantification of C05-05 burden in a 200-pixel (166 µm) radius from a C05-05^-^ neuron. **e-f:** Correlation between mean z-score of calcium fluorescence traces of individual C05-05^-^ neurons and local C05-05 burden within a radius of 200 pixels (166 µm) from PFF-injected mice under visually evoked (*e*) and spontaneous (*f*) conditions at 4-5 MPI. Pearson correlation coefficient (r = 0.0589, *p* = 1.68 × 10^−6^, n = 6264 neurons in *e* and r = 0.121, *p* = 6.49 × 10^−29^, n = 8470 neurons in *f*).

### Sparse α-syn pathology in V1 does not impair visuoperceptual function

Visuoperceptual impairments are common in PD patients^10^. To directly test how cortical α-syn pathology impacts visuoperceptual function, we implemented a cortex-dependent coherent motion discrimination task using random dot kinematograms (RDKs)^52,72^.

Importantly, this task is translationally relevant as a similar task is sensitive for the detection of visuoperceptual impairments in patients with PDD and DLB^41^. We trained PFF-injected mice and controls on a go/no-go version of this task. Starting ∼5 days after surgery, mice were habituated to handling and head-fixation and water deprived to motivate task performance. Mice were then trained to discriminate RDKs drifting in one of two directions, either licking for a water reward to the 0° (rewarded, “Go”) stimulus or withholding licking to the 270° (unrewarded, “No-Go”) stimulus (**Fig. 6a**). Mice were initially trained at 90% dot coherence, and performance was quantified using a discriminability index (d’). PFF-injected and control mice could both learn the task, with PFF-injected mice exhibiting a trend toward slower learning, though this was not significant (**Fig. 6e**, sessions to criterion of d’ ≥ 1.7, 7.33 ± 0.63 vs. 9.67 ± 0.80 days in control and PFF-injected mice, respectively, *p=*0.795).

**Figure 6:**
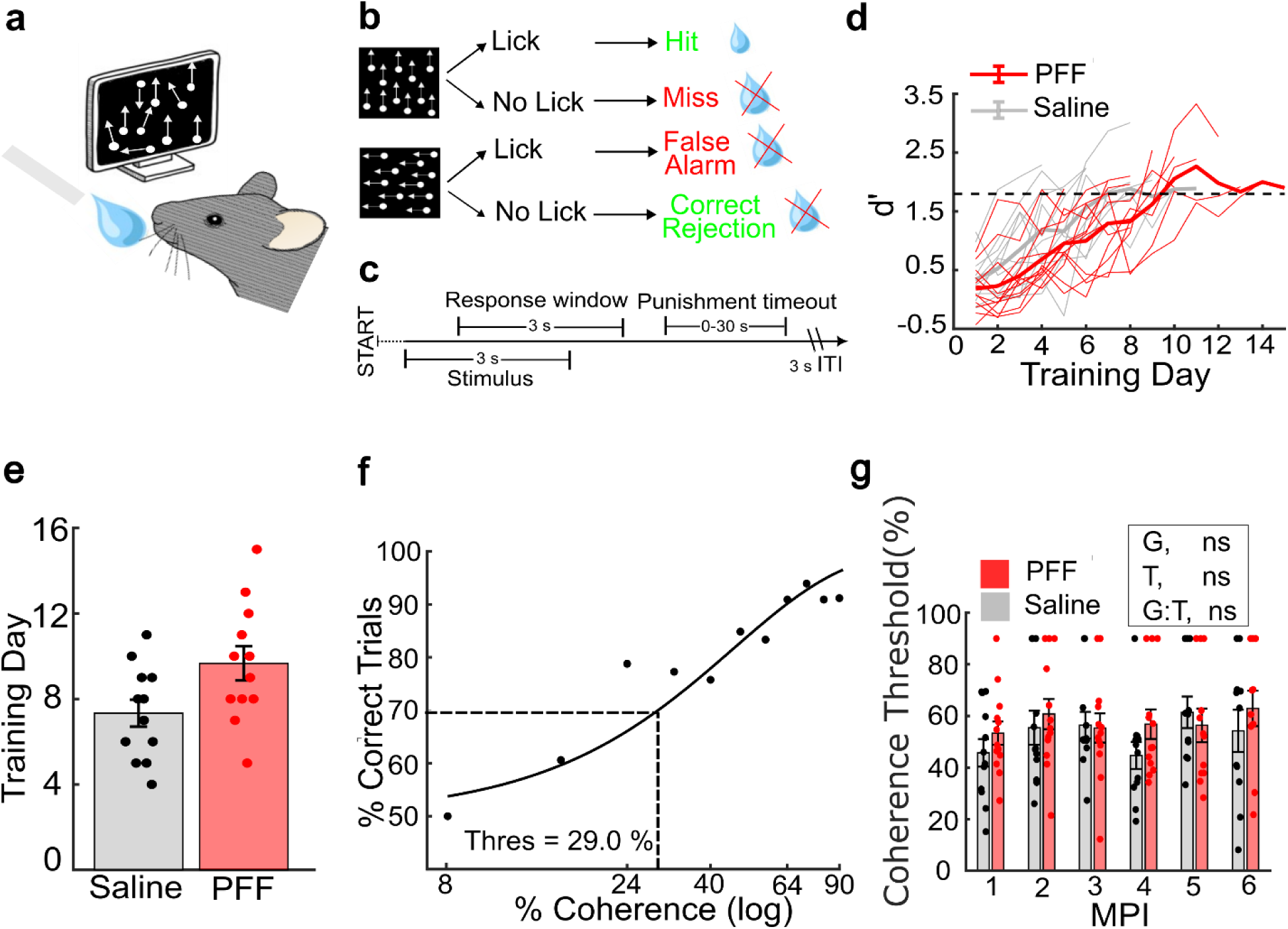
Sparse α-syn pathology in V1 does not impair visuoperceptual function. **a**: Schematic of the Go/No-Go behavioral task, with visual stimuli presented contralateral to the injected hemisphere and lickport for detecting responses and delivering rewards. **b**: Task structure. RDK stimuli moving at 0° (rewarded, “Go”) or 270° (unrewarded, “No-Go”) are presented and trial outcomes based on response are shown. **c**: Timeline of an individual trial for the task. **d**: Task learning of PFF-injected and control mice quantified using discriminability index (d’) across training sessions. Thick dark lines depict the group means with thin, lighter lines for individual mice. Horizontal dotted line shows learning criterion, d’ = 1.7. **e**: Mean training sessions to reach learning criterion (Two-sided Wilcoxon rank sum test, *p* = 0.795. n= 12/group). **f**: Performance of one control mouse across test coherence values, with Weibull curve fit and coherence threshold (29.0%) shown. **g**: Coherence thresholds over time up to 6 MPI in PFF-injected and control mice. LME model, ANOVA for fixed effects of group (G, *p* = 0.363), timepoint (T, *p* = 0.170), and group-by-timepoint interaction (G:T, *p* = 0.590) were not significant. Bars show mean ± s.e.m., with data points for individual mice (n=12 control and n=13 PFF-injected) overlaid.

After learning the basic task, we introduced test sessions in which we interleaved stimuli with coherence values ranging from 8 to 80% (with 8% spacing) between every 1-3 trials of stimuli at training coherence values (90%). Training trials at 90% coherence served as an internal control for task engagement and motivated continued task participation (see Methods). After obtaining >40 trials at each test coherence value (over 3.85 sessions, on average) Weibull curves were fitted^53^ to performance for each mouse and we computed coherence thresholds, defined as the coherence at which performance drops to 70% correct (**Fig. 6f**). Mice were re-tested monthly up to 6 MPI as α-syn pathology progressed. Performance of individual mice in both groups fluctuated over time (**Fig. S13a-b)**. At the group level, coherence thresholds in controls were stable over time (**Fig. 6g**). PFF-injected mice exhibited a trend toward higher coherence thresholds across most timepoints, but there was no statistically significant group-by-time interaction. Performance of PFF-injected and control mice was similar in both “Go” and “No-Go” trials, suggesting that performance of mice is not biased by trial types (**Fig. S13c-d).** Latency to first lick was qualitatively similar between groups and across coherence thresholds (**Fig. S13e-f).** This data suggests that visuoperceptual function is largely preserved in mice with sparse α-syn pathology in V1, at least until 6 MPI.

## Discussion

Cognitive impairment is common in patients with PD and is a defining feature of DLB^2,4,5,7^. However, the mechanisms underlying these cognitive impairments remain largely unknown. The aggregation of α-syn into intraneuronal Lewy-pathology is a hallmark of both diseases and has been widely hypothesized to play a causative role. Across populations, Lewy pathology in specific brain regions correlates with specific symptoms^24,28^. In the case of cognitive impairment, the presence and extent of α-syn pathology in limbic and neocortical regions strongly correlates with symptoms^25,26^. Yet, there is marked inter-individual variability between the burden of neocortical α-syn pathology and extent of cognitive impairment, and there is little direct evidence as to whether, or how, α-syn pathology affects cortical function. To address this question, we developed a model in which we could seed α-syn pathology directly in a disease relevant cortical region. We focused on V1 as posterior cortical impairments are correlated with progression of cognitive impairment and development of dementia in persons with PD^9,11,17,20,41^. Using mouse PFFs injected directly into V1, we observed the development of phosphorylated-S129+ Lewy-like α-syn pathology that evolved over time. The density of pathology increased over time in most regions (**Fig. 1**), with a time course similar to the time course of pathology observed after PFF injection into the dorsolateral striatum^60,61^. Even at 3 MPI, pathology was relatively sparsely distributed. While few studies have carried out detailed neuropathological quantifications in visual areas, sparse α-syn pathology is commonly observed in primary and higher order visual areas in persons with PD and DLB^62–64^. Thus, our model achieves translationally relevant levels of α-syn pathology in V1.

Within V1, α-syn pathology showed a clear laminar distribution with a preference for deeper layers (L5, and L6a) (**Fig. 1**). Cortical layer-specific seeding has been best characterized in the motor cortex after striatal injection, where pathology is most abundant in L5 and L2/3^39,73,74^ and is thought to reflect retrograde transmission^61^. Since we injected directly in V1, our data adds to other studies suggesting that there may be layer- or cell-intrinsic factors that increase vulnerability to the uptake of PFFs and/or the formation of α-syn pathology^55^. We also observed α-syn pathology develop over time in brain regions distant from the injection site, including the contralateral V1, dorsomedial striatum, secondary motor and prefrontal cortical areas, and hippocampus (**Fig. 1**). Most of these brain regions either receive projections from or project directly to V1^75,76^, suggesting spread through anatomical connections as has been well documented for other PFF injection sites^61,77,78^. Interestingly, α-syn pathology was present in the dorsomedial striatum, which receives a direct projection from V1, but not the dorsolateral geniculate nucleus, which projects to V1. This suggests that the dLGN is relatively resilient to α-syn transmission, consistent with other prior studies showing resilience to formation of α-syn pathology in the thalamus^79,80^. Alternatively, anterograde transmission may predominate from V1, in contrast to the primarily retrograde transmission that occurs from the dorsolateral striatum^61^.

Having established a model with posterior cortical α-syn pathology, we next investigated how this pathology affects visual cortical circuit function. At the population level we observed small, but significant changes, with a shift toward more visually responsive neurons with lower orientation and direction selectivity in PFF-injected mice compared to controls (**Fig. 2f-j**). These changes evolved over time and were most prominent at 4-5 MPI, time points that correspond with peak α-syn pathology. There were also changes in spontaneous activity during quiet wakefulness across the population of all recorded neurons, including an increase in the frequency of calcium transients and a slight reduction in transient amplitude at 4-5 MPI (**Fig. 3c-d**). Although these results are statistically rigorous, using a large sample size and mixed methods approaches, the absolute magnitude of these effects is small and of uncertain biological significance. Our behavioral results suggest that such small circuit-level changes do not functionally impair visuoperceptual performance, as discussed further below.

To investigate whether cells with somatic α-syn inclusions might be differentially affected, we took advantage of C05-05, a recently characterized fluorescent ligand that binds specifically to aggregated α-syn^40^. Compared to neighboring C05-05^-^ neurons, neurons with large somatic α-syn inclusions showed significant reductions in visually evoked activity across a range of activity parameters (**Fig. 4c-f**). Interestingly, we also observed a correlation between the density of the somatic C05-05 signal and reductions in neuronal activity (**Fig. 4h-i**). In other words, neurons with the most substantial intracellular α-syn burdens and mature somatic inclusions showed the greatest reductions in neuronal activity. Conversely, neurons without inclusions showed higher neuronal activity that correlated with the surrounding burden of α-syn pathology (**Fig. 5b-c, e-f**). Functionally, visuoperceptual ability was largely preserved in PFF-injected mice compared to controls, even up to 6 MPI (**Fig. 6e**).

Together, our data suggest a model in which sparse α-syn pathology drives reductions in activity in neurons with large, somatic intracellular inclusions. This effect could be directly due to toxic effects of α-syn on the neuron bearing the inclusion. Alternatively, α-syn has important roles in presynaptic function and neurotransmitter release^32,33,81^ and reductions in activity could be a result of reduced synaptic inputs, though we would not necessarily expect reduced activity to be specific to inclusion-bearing neurons in this case. Neuroinflammatory changes could contribute as well, though we do not see signs of overt neuroinflammation in our model (**Fig. S3-S4**). The increased activity seen in inclusion-free neurons may represent a homeostatic network response to hypofunctional inclusion-bearing neurons. This interpretation could explain the changes we observed in population activity in PFF-injected mice compared to controls: Increased activity in inclusion-free neurons resulting in more visually responsive neurons with lower orientation and direction selectivity. Similar network changes have been observed in mouse models with other neurodegeneration-associated pathologies. For example, in mice with extracellular Aβ plaques, orientation selectivity of neurons in V1 decreases over time as pathology develops, with an increase in both hypo- and hyperactive neurons^82,83^. On the other hand, in mice overexpressing human P301L tau, cortical neurons are hypoactive, but this phenotype is driven by soluble tau, with inclusion-bearing neurons showing normal visual evoked activity^71,84^. Therefore, there may be convergent and divergent mechanisms of circuit dysfunction across different neurodegenerative diseases, and it remains important to investigate the effects of these neurodegeneration-associated pathologies independently and as co-pathologies.

Our study adds to emerging evidence on the effects of intraneuronal accumulation of α-syn on neuronal activity. A few studies have investigated the effects of α-syn pathology in vitro, with mixed results. In rat cortical neuron cultures, treatment with recombinant human α-syn oligomers led to a time-dependent reduction in neuronal activity^33^. In mouse hippocampal neuron cultures, treatment with PFFs has been reported to both increase^32^ and decrease^85^ miniature excitatory post-synaptic potentials, and to decrease somatic calcium transients^32^. In vivo, even fewer studies have investigated the effects of α-syn on cortical circuit function and neuronal activity. After injection of PFFs into the dorsolateral striatum, Blumenstock et al. found a progressive loss of dendritic spines^36^ and increased spontaneous calcium transients and greater responses to whisking at 9 MPI^38^ in the somatosensory cortex (S1). Recently, Khan et al. reported very large increases in locomotion induced firing rates in neurons from primary motor cortex (M1) after injection with PFFs in either M1 or dorsolateral striatum^39^. Given that these studies could not differentiate neurons with and without intraneuronal α-syn inclusions, it is possible that these observations of increased activity in M1 and S1 reflect a shared homeostatic response to hypofunctional inclusion-bearing neurons. Consistent with α-syn inclusions causing reduced neuronal activity, using a novel *SCNA* fluorescent reporter knock-in mouse line, Zhang et al. recently reported hypoactivity of inclusion-bearing neurons in M1 after PFF injection into M1^86^.

The mechanism(s) underlying effects of α-syn inclusions on neuronal activity remain incompletely understood. Intracellular pathology progresses from neurites to the soma, with initial α-syn aggregation enriched in pre-synaptic terminals and axons^35,44,87^, and leading to reduced excitatory synaptic neurotransmission^88^. As pathology then progressively involves dendrites and finally the soma, dendritic spines are lost resulting in reduced excitatory inputs onto inclusion-bearing neurons^32,36,73,89^. Aggregation of α-syn has also been reported to affect the intrinsic excitability of neurons. In ex vivo slices after intrastriatal PFF injections reduced excitability has been observed in striatal spiny projection neurons^90^ whereas increased excitability was found in neurons in the secondary motor cortex^73^. Thus, it is possible that there are regional- or cell-type-specific effects of α-syn pathology on neuronal activity. Future studies comparing neurons with and without inclusions across multiple brain regions and neuronal sub-types will be required to more comprehensively understand how α-syn pathology impacts circuit function.

Our study has important limitations. Transmission of α-syn pathology is hypothesized to be a key pathophysiological mechanism underlying disease progression in PD^91^. After injecting PFFs in V1 we observed spread to multiple anatomically connected brain regions, reflecting a potentially important translational aspect of the model. However, we cannot be certain that effects we observed in V1 are due exclusively to α-syn pathology in V1. It is probable that α-syn pathology also affected neuronal activity in other brain regions, especially those to which there was significant spread. This could have affected activity in V1 through long-range projections to V1 or feedback mechanisms. To measure neuronal activity, we used jRGECO1a, a red-shifted calcium indicator.

Although calcium indicators are widely used as a proxy for neuronal activity, changes in intracellular calcium handling induced by α-syn pathology could potentially confound our interpretations. For example, changes in intracellular buffering of calcium have been reported in a transgenic mouse overexpressing human α-syn, leading to elevated calcium transients in the absence of alterations in spiking activity^37^. Whether impaired buffering of intracellular calcium happens in neurons in V1 in the PFF model is unknown, but if present, this effect might have led us to underestimate the impact of α-syn inclusions on neuronal activity. We focused on imaging with C05-05 at 4–5 MPI as this represents the peak of α-syn pathology in L2/3 of V1. Imaging the effects of α-syn pathology on neuronal activity at earlier and later timepoints would also have added valuable information on the time-course of the observed effects. However, at earlier time points (1-3 MPI) we observed minimal somatic α-syn pathology and at later timepoints (6 MPI) there was a decline in somatic α-syn pathology observed in vivo, possibly reflecting degeneration of inclusion-bearing neurons^35^. C05-05 signals were difficult to resolve beyond L2/3. Sparse in general, α-syn pathology was less common in L2/3 compared to deeper layers. It is possible that the effects we observed, particularly at the population level, were small for this reason. Future studies should investigate deeper layers, with more abundant pathology, where changes in neuronal activity may be more robust. Given the reductions in neuronal activity in inclusion-bearing neurons and the necessity for V1 in apparent motion discrimination^52^, we speculate that if the burden of pathological α-syn were substantially increased in our model, behavioral impairments may emerge.

The use of C05-05 to differentiate neurons with and without somatic inclusions facilitated key findings in our study. However, it is likely that many neurons had neuritic, but not somatic, α-syn aggregates. Indeed, we observed abundant, small C05-05 signals that did not localize to cell somas (**Fig. 4a,g**). It was impossible to trace these signals back to individual cell bodies and thus, we could not determine how isolated neuritic α-syn pathology might affect neuronal activity. By manually selecting neurons with somatic α-syn inclusions, there is the possibility for some unconscious bias in our results. To verify our findings, we used an automated neuron selection method and importantly, found a clear relationship between the density of intraneuronal α-syn pathology and reduced neuronal activity (**Fig. 4h-i**). As this automated method relied on a functional algorithm to detect active neurons with jRGECO1a calcium transients, very hypofunctional, inactive neurons were likely missed. Indeed, we observed many large, somatic appearing α-syn inclusions that were not co-localized with jRGECO1a auto-segmented somas (**Fig. 4g**). So, we believe our automated analysis is likely underestimating the effect of α-syn pathology on inclusion-bearing neurons, if anything.

Our study has identified important neuron and circuit level effects of α-syn pathology. Inducing sparse pathology in V1, our model is consistent with early-stage neocortical spread of α-syn pathology seen in patients with PD and DLB. Our results show, for the first time in vivo, that somatic α-syn inclusions lead to hypoactivity of posterior cortical neurons. The increase in neuronal activity in surrounding cells without inclusions may be compensatory, helping to preserve visuoperceptual function in the face of accumulating α-syn pathology, as observed in our PFF-injected mice with preserved visuoperceptual function. Indeed, reduced performance on a coherent motion task was seen only in PD patients with dementia, and not PD patients with normal cognition^41^, suggesting decline in visuoperceptual ability represents a functional transition in the progression of PD-related cognitive impairments. Our results also suggest a framework to understand inter-individual variability in the relationship between α-syn pathology and cognitive impairments amongst patients. Patients with preserved network plasticity mechanisms may be able to compensate for some α-syn pathology and maintain cognitive function. In the future, this model can serve as a foundation for understanding how risk factors for dementia in PD, including aging, other neurodegenerative co-pathologies, and genetic risk factors, interact with α-syn pathology to drive cognitive impairment.

## Methods

### Experimental animals

All experiments followed the US National Institutes of Health guidelines for animal research, under an animal use protocol approved by the University of California, Los Angeles Animal Research Committee. Male and female mice were used, beginning at 7–12 weeks of age at the time of cranial window surgery or PFF/saline injections. All animals were housed in a vivarium with a reverse 12 h light/dark cycle in groups of 3–5 animals per cage. For immunohistochemistry staining to characterize PFF induced pathology, we used wild-type C57BL/6J mice (strain #000664, The Jackson Laboratory). For in vivo imaging and motion discrimination behavioral tasks, we used Thy1-jRGECO1a transgenic mice^42^ (strain #030526, The Jackson Laboratory) which were maintained on a C57BL/6J background.

### In vitro α-syn PFF generation

Full-length mouse α-syn (1-140) monomers were expressed in BL21 (DE3) RIL cells and purified according to previous protocols^43^. Mouse PFFs were generated by diluting purified α-syn monomers to 5 mg/ml in sterile Dulbecco’s PBS (Cellgro, Mediatech; pH adjusted to 7.0) and then incubating at 37°C with constant agitation at 1,000 r.p.m. for 7 days. The sedimentation test and thioflavin T-binding assay were used to verify α-syn fibrillization, as previously described^44^. The seeding potency of the sonicated PFFs was verified using a mouse primary cortical neuron culture assay. To confirm the formation of true pathological aggregates, we treated the cells with 1% Triton X-100 (TX) prior to fixation to remove soluble proteins, then performed immunofluorescence staining for α-syn aggregates using an anti-p-S129 α-syn antibody (BioLegend, #825701) (**Fig. S14**). The presence of these robust, Triton X-100-insoluble puncta confirmed that the sonicated PFFs possessed high seeding activity.

### PFF injection and Cranial Window Surgery

Mice were deeply anesthetized using 5% isoflurane followed by maintenance with 1.5–2% isoflurane. For calcium imaging and motion discrimination tasks, cranial window implantation^45,46^ and PFF stereotaxic injection was performed in the same surgical procedure. After removing the scalp and periosteum, a ∼3-mm-diameter circular craniotomy, centered ∼2.5 mm lateral to the midline and ∼0.5 mm anterior to the lambda suture was created using a pneumatic dental drill with an FG 1/4 drill bit (Kerr Dental) over the left V1. Using a microinjector (Neurostar) with a 33 g needle (Hamilton, 80408), 2.5 µL of freshly sonicated PFFs (2.5 µg/µl) were injected into each of three sites across V1 (0.5 mm anterior to lambda and 2 to 2.5 mm lateral to the central suture) at a rate of 200 nL/min followed by a 5 min waiting period to prevent backflow. A 3 mm coverslip glued to a 5 mm coverslip using an optical adhesive (Norland Products, catalog #71) was then placed over the craniotomy. The coverslip was gently pressed against the brain and was glued in place to the skull with cyanoacrylate glue (Krazy Glue) followed by black dental cement (OrthoJet, Lang Dental). A customized stainless-steel headbar was then attached to the skull using dental cement to allow subsequent fixation of the mouse onto the microscope stage. For immunohistochemistry staining to characterize PFF pathology, after PFF injection, the 3-mm craniotomy was sealed with a coverslip, and the skull was covered by dental cement. Carprofen (5 mg/kg, i.p.; Zoetis) and dexamethasone (0.2 mg/kg, i.p.; Vet One) were injected subcutaneously for pain relief and mitigation of edema on the day of surgery and daily for the next 48 h.

### Intrinsic optical signal imaging

IOSI was performed as previously described^47^. Animals were sedated with chlorprothixene (∼3 mg/kg, i.p.), lightly anesthetized with ∼0.5-0.7% isoflurane, and head-fixed under the microscope. The cortical surface was illuminated with 525 nm light to capture an image of the superficial vasculature. The microscope was then focused 300 µm below the cortical surface and illuminated with 625 nm light to record intrinsic signals. Frames were collected at 10 Hz (100 ms exposure time) using an 8-megapixel CCD camera (Thorlabs, 8051M-USB) during 30 trials of a series of standard drifting sinusoidal gratings with 1.5 seconds duration. Visual stimuli were generated with custom scripts based on PsychoPy^48^. The cortical V1 map was generated as previously described^47^. Briefly, stimulus-evoked change in reflectance values (ΔR/R) were calculated and binarized by thresholding for ΔR/R values below a Z-score of −3. Binarized images were then pseudocolored and overlaid onto images of the vasculature.

### Two-photon imaging

In vivo imaging was performed using a Thorlabs Bergamo II microscope with fast galvo-resonant scanning mirrors, 14° collection optics with two high-sensitivity GaAsP photomultiplier tubes with green (525/50 nm) and red (607/70 nm) emission bandpass filters, and a 16x/0.8 NA objective (Nikon), coupled to an Insight X3 dual output Ti:Sapphire laser (Spectra Physics). The fixed 1045 nm line was used to excite jRGECO1a. During the recording periods animals were awake and stationary sitting in a cylindrical tube for comfort. The animals were habituated to head-fixation in the tube prior to imaging. Time-series movies of jRGECO1a fluorescence were acquired with bidirectional scanning at a frame rate of ∼10 Hz (512 x 512 pixels field of view; 0.825 µm/pixel). One to two focal planes from depths of around 200-250 µm were acquired in each mouse at all timepoints. We also acquired more superficial imaging planes (50-150 µm) where C05-05 signals were more common at 4-6 MPI. Regions of interest in V1 were chosen according to IOSI maps. Visual stimuli were generated with custom scripts using PsychoPy^48^. A monitor (DELL P2219H LED, with 1920 × 1080 pixels, 21.5” measured diagonally, 60 Hz refresh rate) was positioned 15 cm from the mouse’s right eye (contralateral to the craniotomy). A series of standard drifting sinusoidal gratings (8 directions, 0 to 315° in 45° increments) with a spatial frequency of 0.04 cycles per degree and a temporal frequency of 1 Hz were displayed in a pseudorandom order for 1 second with 3 second inter-stimulus intervals (gray screen) between the stimuli. Each of the 8 directions were presented for four trials over 150 seconds during the imaging session. Visual stimuli timing data was captured with a photodiode (Bpod Frame2TTL, Sanworks) and synchronized with 2P frame acquisition times using ThorSync software. To reduce light contamination, the conical tip of the objective was shielded from light coming from the screen by removable mounting putty. We also recorded spontaneous activity with mice head fixed in front of a black screen for ∼300 seconds. Imaging was performed monthly after PFF injection for up to 6 months. Over time, mice were lost due to window opacification or bone growth (Table 1). The total number of neurons recorded, and the mean number of neurons recorded per field of view for visually evoked and spontaneous activity are described in Tables 2 and 3.

**Table 1.** Number of mice used for in vivo imaging.

| MPI | 1 | 2 | 3 | 4 | 5 | 6 |
| --- | --- | --- | --- | --- | --- | --- |
| PFF | 22 | 24 | 23 | 15 | 16 | 12 |
| Saline | 19 | 19 | 17 | 14 | 16 | 12 |

**Table 2.**
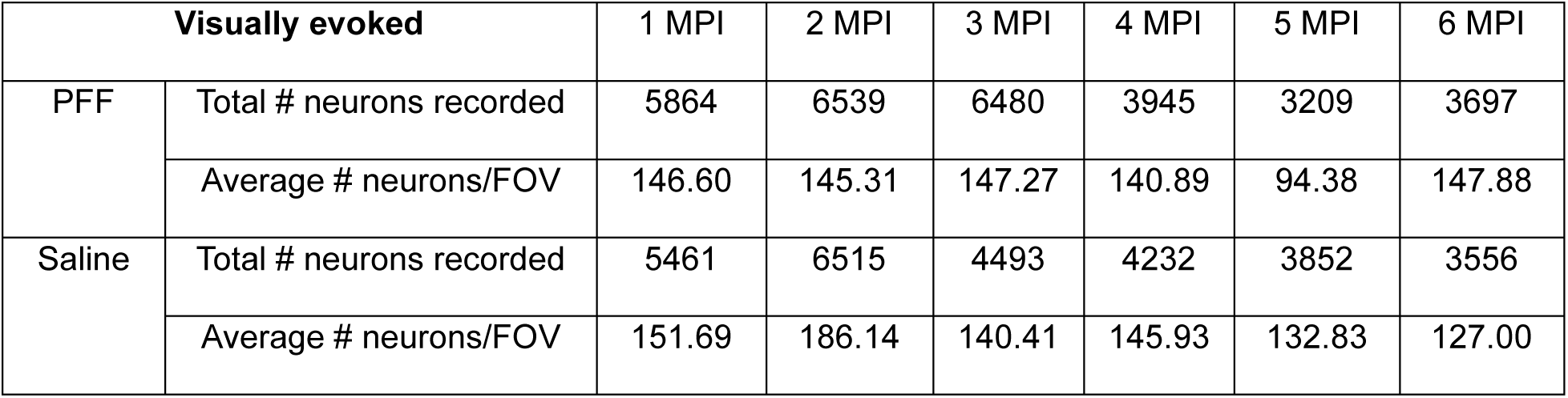
Number of neurons recorded in visually evoked conditions.

| Visually evoked |  | 1 MPI | 2 MPI | 3 MPI | 4 MPI | 5 MPI | 6 MPI |
| --- | --- | --- | --- | --- | --- | --- | --- |
| PFF | Total # neurons recorded | 5864 | 6539 | 6480 | 3945 | 3209 | 3697 |
|  | Average # neurons/FOV | 146.60 | 145.31 | 147.27 | 140.89 | 94.38 | 147.88 |
| Saline | Total # neurons recorded | 5461 | 6515 | 4493 | 4232 | 3852 | 3556 |
|  | Average # neurons/FOV | 151.69 | 186.14 | 140.41 | 145.93 | 132.83 | 127.00 |

**Table 3.** Number of neurons recorded in spontaneous conditions.

| Spontaneous |  | 1 MPI | 2 MPI | 3 MPI | 4 MPI | 5 MPI | 6 MPI |
| --- | --- | --- | --- | --- | --- | --- | --- |
| PFF | Total # neurons recorded | 9076 | 9873 | 10139 | 5897 | 4780 | 5294 |
|  | Average # neurons/FOV | 221.37 | 210.06 | 206.92 | 210.61 | 144.85 | 196.07 |
| Saline | Total # neurons recorded | 8257 | 9390 | 7827 | 6432 | 6023 | 4974 |
|  | Average # neurons/FOV | 235.91 | 268.29 | 217.42 | 207.48 | 207.69 | 177.64 |

### C05-05 Imaging

We used C05-05^40^ to visualize α-syn aggregates. At 4 and 5 MPI, 4 µl/g of C05-05 (4 mg/ml in dimethyl sulfoxide, total 1.6 mg/kg) was injected intraperitoneally 90 minutes before 2P imaging. C05-05 was excited at 800 nm and emitted fluorescence was captured in the GFP channel. Aggregated α-syn was visualized only in mice injected with PFFs. Calcium imaging of spontaneous and visually evoked activity was recorded in the same FOVs where α-syn aggregates were concentrated.

## 2P Imaging Analysis

Motion correction, image segmentation and calcium signal extraction were performed using the python 3.6 package Suite2P^49^. Non-rigid motion-correction was performed using default settings. Region of interest (ROI) detection was performed with the following parameter settings: tau (0.7), roidetect (1), sparse mode (1), denoise (1), spatial scale (1), connected (1), threshold scaling (1), max overlap (0.75), max iterations (20), high pass (100) and spatial hp detect (25). Additional data analysis was performed in MATLAB. Neuropil extraction was performed by subtracting 0.7 times the neuropil fluorescence signal from somatic fluorescence signals. We calculated a modified Z score vector (Zf) from the fluorescence traces for each neuron as previously described^50,51^. We defined cells as active if the cell had at least 0.5 seconds (five consecutive frames) with Zf > 3 during the recording. To classify individual cells as visually responsive, we determined if cells showed time-locked responses to any visual stimulus using a probabilistic bootstrapping method as described previously^50^. Briefly, we calculated the correlation between the trace of all visual stimuli (stimulus on = 1, stimulus off = 0) and the Zf vector from each individual cell. We then created 1,000 scrambles of Zf, preserving calcium transient epochs (epoch ≥ five consecutive frames with Zf ≥ 3), calculated correlations between the stimulus trace and each scrambled calcium trace, and generated a distribution of correlation values for each of the 1000 scrambled traces. If the true correlation for a cell was in the 99^th^ percentile of the distribution of scrambled correlation values (*p* values < 0.01), then we considered that cell as being visually responsive.

Next, modified Z-scores for each of the 32 individual visual stimuli were aligned to the onset of the visual stimulus and averaged across the four trials displayed for each stimulus direction. The area-under-the-curve (AUC) of the mean stimulus evoked trace for each stimulus direction was quantified as the trapezoidal integral of the modified Z-score vector over 1.5 s after stimulus onset. The preferred direction and orientation were identified as the stimulus direction or orientation evoking the largest AUC. The orientation selectivity index (OSI) and direction selectivity index (DSI) of each cell was calculated as:

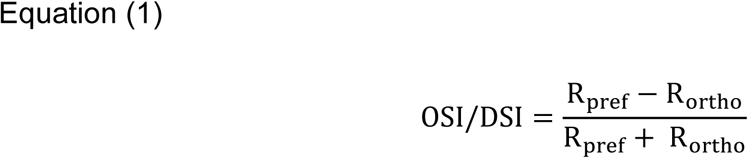

where *R_pref_* is the AUC of the mean response to the preferred orientation/direction and *R_ortho_* is the AUC of the mean response to the orthogonal orientation/direction. OSI/DSI values thus spanned 0 to 1, with an equal response to all orientations/directions having OSI/DSI = 0, and perfectly selective responses to only one orientation/direction having an OSI/DSI = 1. Highly orientation or direction selective neurons were defined as neurons with an OSI or DSI greater than 0.6. For each neuron we also calculated characteristics of visually evoked calcium transients during individual stimulation epochs using the “findpeaks” function in MATLAB, with settings of “MinPeakHeight” = 4 and “MinPeakDistance” = 4 frames. For spontaneous activity, amplitude and frequency of calcium transients were detected across the entire ∼ 300 second calcium trace using the “findpeaks” function in MATLAB with the same settings. We also calculated the mean Zf and total AUC of the entire trace. Modified Z-scores for each of the 4 individual visual stimuli of preferred orientation were aligned to the onset of the stimulus and a mean stimulus evoked trace was calculated for each cell. A modulation index (MI) indicating the strength of the visually evoked response for each cell was calculated as:

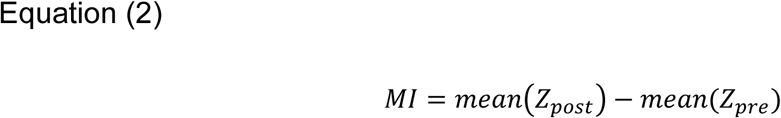

where *mean(Z_post_)* and *mean(Z_pre_)* are equal to the mean Zf for 2 seconds after and before stimulus onset, respectively.

We analyzed pairwise synchronicity of cells and ensemble detection as previously described^39^. Briefly, calcium spike raster matrices were generated from neuropil subtracted change in fluorescence signals (ΔF/F) by thresholding the first derivative at 3 standard deviations above the baseline fluorescence activity. SØrenson-Dice correlation (SDC) of spike matrices from visually evoked and spontaneous conditions were then calculated as previously described^39^. To calculate average pairwise synchronicity across timepoints, we applied a bootstrapping approach in which each cell’s spike train was circularly shifted by a random time offset ranging from 1 to the number of frames of a video. This process was repeated 1,000 times for all cells. Pairwise connections were considered significant if their observed synchrony exceeded the 99th percentile of the shuffled distribution. Ensemble detection in the primary visual cortex was performed as previously described^39^. The similarity index between cells was first calculated to define ensembles in which cells participated under visually evoked and spontaneous conditions. Ensemble stability was then defined as the proportion of neurons in a given ensemble that participated in the ensemble during its reactivation.

To manually select individual jRGECO1a^+^ cells with somatic α-syn inclusions, we used raw images of C05-05 signals and merged these with single-plane images of jRGECO1a^+^ fluorescence, temporally averaging 20-30 frames to improve signal-to-noise ratio. Using Suite2P, we selected cells in which C05-05 signals clearly and completely overlapped with jRGECO1a^+^ somas and defined them as C05-05^+^ cells. We then defined C05-05^-^ cells by selecting neighboring jRGECO1a^+^cells without C05-05 signals within ∼200 µm from each C05-05^+^ cell. To quantify C05-05 signals, we binarized C05-05 images in MATLAB by adaptive thresholding using the function “imbinarize” and manually adjusting the sensitivity factor for each image. The binarized images were then median filtered (“medfilt2”) by setting a 5 x 5 window around a pixel to be smoothed. Binarized C05-05 signals were then overlaid onto jRGECO1a^+^ soma coordinates segmented by Suite2p. We calculated the density of C05-05 in a cell by dividing the total number of C05-05 pixels within a cell by the total number of pixels segmented for that cell in Suite2p and excluded cells with very low C05-05 burdens (less than 20 C05-05 pixels) which tended to represent background noise. To correlate total α-syn burden with local neuronal activity, we summed up the total binarized C05-05 signal in each FOV and correlated this with neuronal activity measures from jRGECO1a^+^ /C05-05^-^ cells in the corresponding FOV. To correlate local α-syn burden with the activity of individual cells, we summed the total binarized C05-05 signals in a circle circumscribed by a defined radius (e.g. 200 pixels, ∼166 µm) from the centroid of C05-05^-^ cells and divided that by the area of the region to obtain the local density of C05-05 signals.

### Go/No-Go visual motion discrimination task for head-restrained mice

A custom-built behavior testing rig was used to assess visual motion perception in mice. The behavioral task was controlled and synchronized using a Bpod State Machine r2 system (Sanworks). A custom made lickport connected to a capacitive touch sensor (Sparkfun AT42QT1011) was used to monitor licking behavior. Mice were head fixed in a cylindrical tube on a platform, with a monitor (DELL P2225H LED, with 1920 × 1080 pixels, 21.5” measured diagonally, 60 Hz refresh rate) positioned at a distance of 15 cm from the right eye of the mouse (contralateral to the PFF-injected hemisphere). Visual stimuli were generated using the ‘Psychtoolbox’ package in MATLAB (release 2022b; Mathworks). We adapted random dot kinematogram (RDK) stimuli for use in mice as described previously^52^. White dots on a black background were generated randomly and uniformly to cover the entire screen of the monitor. All dots within a kinematogram shared the same size (2.37°), lifetime (100 frames), and step size (800 pixels per second). For each trial the number of coherently moving dots was set between 8 and 90%, with the remaining dots moving in random directions selected uniformly across all possible angles. Dot group assignments remained fixed throughout the stimulus duration. Dot motion was set to 0° (up) for rewarded “Go” trials and to 270° (left) for unrewarded “No-Go” trials.

Approximately 5 days after PFF injection surgery, mice were subjected to handling for 10 minutes daily until they were comfortable with the experimenter (typically 3-5 days). Mice were then habituated to head-fixation in a behavioral enclosure for 10 minutes on two consecutive days. Animals were monitored during behavior with a webcam and behavior was terminated early if there were any signs of distress observed. Concurrently, mice were water restricted with ad libitum access to food. Body weight was checked daily. On days in which mice were habituated or behavioral training did not occur, mice were given ∼0.5-1 ml of water to achieve a goal of ∼15-20% weight loss, which was maintained during training. Mice with weight loss >25% or signs of ill health were removed from water restriction. Mice were first trained to lick to trigger water rewards (∼5 µl). Once mice learned to lick, we introduced “Go” RDK stimuli at 90% coherence and each stimulus, presented for 3 seconds, was coupled with a water reward dispensed in response to licking occurring at least 1 s after stimulus presentation. Once mice learned to lick in response to the stimulus and withhold licking in the intertrial interval ranging from 3 to 10 seconds (licking on ∼80% of trials in a session for 1-2 days), mice were advanced to the Go/No-Go task. Mice were trained to discriminate between “Go” and “No-Go” stimuli presented in random order, with no more than 3 trials of the same stimulus type occurring consecutively. Licking to the “Go” or “No-Go” stimuli was classified as a “Hit” or “False Alarm” (FA), respectively. Withholding licking to the “Go” or “No-Go” stimuli was classified as a “Miss” or “Correct Rejection” (CR), respectively. FA responses triggered an additional punishment time ranging from 3 to 30 seconds, gradually increased by the experimenter, after the intertrial interval to discourage impulsive licking.

Performance of the mouse was quantified using a discriminability index (d’), calculated as:

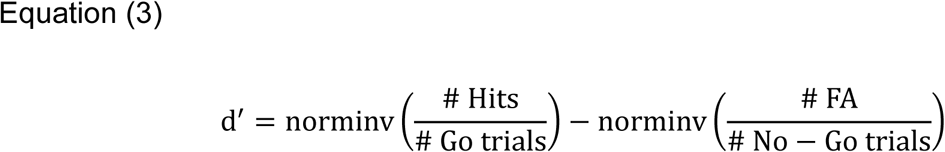

where norminv is the inverse of the normal cumulative distribution function. Mice were trained until achieving a d’ ≥ 1.7 for 1-2 days. To determine motion discrimination coherence thresholds, test stimuli with coherence values ranging from 8 to 80% (with 8% spacing) were introduced between trials of stimuli at training coherence values (90%). If performance on training trials across a session dropped below d’ = 1.8 or impulsive licking was observed, the ratio of training to test trials was set at 3:1; otherwise, we used a ratio of 1:1. Both testing and training trials had the same punishment time and intertrial intervals. Testing sessions lasted for 237.4 trials on average and sessions were continued until mice had completed 40 trials per test coherence value (average = 3.85 sessions). The percentages of mean correct trials for each coherence were fit with a Weibull cumulative distribution function in MATLAB^53,54^. The slope (β) and threshold parameters (α) were obtained by optimizing the log likelihood comparing the observed psychometric trial data to the data generated by the Weibull function, and coherence threshold for performance was considered 70% correct. After completing tests for each timepoint, mice were permitted to drink water ad-libitum for about 2 weeks, then placed under water restriction again, re-trained to perform the Go/No-Go task (1-7 sessions to achieve d’>1.8) and were re-tested across all coherence values as described above. Behavioral testing was performed monthly after PFF injection for up to 6 months. Two mice in the PFF group and one mouse in the control group were prematurely euthanized due to signs of ill health (**Table 4**). Pre-training data for one PFF-injected mouse was lost and was therefore excluded from the analysis for pre-training in Fig.6d-e.

**Table 4.** Number of mice used for behavior testing.

|  |  |  |  |  |  |  |
| --- | --- | --- | --- | --- | --- | --- |
| MPI | 1 | 2 | 3 | 4 | 5 | 6 |
| PFF | 13 | 13 | 13 | 13 | 11 | 11 |
| Saline | 12 | 12 | 12 | 12 | 11 | 11 |

### Calcium imaging in primary cortical neuron cultures

Primary cortical neurons were prepared from CD1 wild-type mice and cultured as previously described with minor modifications^55^. Briefly, dissociated cortical neurons were plated in 96-well plates at 25,000 cells per well and maintained under standard neuronal culture conditions. On DIV4-5, neurons were transduced with AAV1-pAAV.Syn.NES-jRGECO1a.WPRE.SV40 (Addgene viral prep #100854-AAV1), at 1 × 10^8^ viral genomes per well. On DIV8, neurons were treated with PFFs at 100 ng per well. Before treatment, PFFs were sonicated in a water bath sonicator (Diagenode) for 20 cycles (30 s on, 30 s off) at approximately 8 °C. On DIV17, spontaneous calcium activity of primary cortical neurons was recorded for 90 s at 10 Hz to obtain baseline measurements. Following baseline recording, C05-05 was added to each well at a final concentration of 100 nM. Neuronal calcium activity was recorded again 30 min after treatment using the same imaging parameters. jRGECO1a and C05-05 were excited at 555 and 405 nm and captured at 615.0 and 530.5 nm respectively using a Nikon microscope (ECLIPSE Ti2. Objective: 20x). Immediately after recording, neurons were fixed with 4% paraformaldehyde (PFA) and permeabilized with 1% Triton X-100.

Neurons were blocked and incubated with primary monoclonal antibody against anti-phospho-α-syn Ser129, 81A (1:4000; BioLegend, #825701) and primary polyclonal antibody against anti-MAP2 (1:4000; BioLegend, #822501). Cultures were then incubated with Alexa Fluor 488 anti-mouse IgG2a antibody (1:2000; Invitrogen, #A-21131) for 81A and Alexa Fluor 647 anti-chicken IgY antibody (1:2000; Invitrogen, #A-21449). Stained neurons were imaged for post hoc assessment of α-syn pathology and neuronal morphology.

To analyze the activity of jRGECO1a^+^ cultured neurons before and after treatment with C05-05, we used the python 3.6 package Suite2P^49^ to perform motion correction, image segmentation and calcium signal extraction. Non-rigid motion-correction was performed using default settings. Region of interest (ROI) detection was performed with the following parameter settings: tau (0.7), anatomical_only (3), diameter (20), cellprob_threshold (0), flow_threshold (1.5), pretrained_model (cyto) and spatial_hp_cp (0). Additional data analysis was performed in MATLAB. We calculated a modified Z score vector (Zf) from the fluorescence traces for each neuron as previously described^50,51^. We then manually selected jRGECO1a^+^ neurons which are both C05-05^+^ and pS129^+^. For each selected neuron, we calculated mean Zf, total AUC, amplitude and frequency of calcium transients using the “findpeaks” function in MATLAB, with settings of “MinPeakHeight” = 7 across the entire 90 s calcium trace.

### Immunohistochemistry

Mice were euthanized with isoflurane and transcardially perfused with PBS, followed by 4% PFA in PBS. After extraction, brains were post-fixed overnight in 4% PFA at 4°C, then washed and preserved in PBS at 4°C. To evaluate the spread of α-syn pathology, 50 µm thick coronal slices were made using a vibrating microtome (Precisionary Instruments Inc, #VF-510-0Z). Brain sections were washed in PBS and blocked with 0.4% Triton-X-100 (Sigma-Aldrich, #X100-500ML), 3% fetal bovine serum (FBS) (Thermofisher, #10437028) and 3% Bovine Serum Albumin (BSA) (Prometheus, #25529) in PBS for 2 hours. Sections were then incubated with primary antibodies against S129-phosphorylated α-syn (Abcam, #ab51253) (1:1000), CaMKII (DSHB, # BCN32.1.3C5) (1:50), Iba1 (Synaptic Systems, 234 308) (1:1000) and GFAP (Invitrogen, #13-0300) (1:1000) and diluted in blocking solution (0.4% Triton-X-100, 3% FBS and 3% BSA in 1xPBS). Sections were then incubated with the corresponding Alexa Fluor-conjugated secondary antibodies: Alexa Fluor 488 anti-rabbit (1:1000; Invitrogen, #A-11008) for S129-phosphorylated α-syn; Alexa Fluor 488 anti-mouse (1:500; Jackson ImmunoResearch, #115-545-003) for CaMKII; Alexa Fluor 568 anti-rat (1:1000; Invitrogen, #A-11077) for GFAP; and Alexa Fluor 647 anti-guinea pig (1:1000; Invitrogen, #A-21241) for Iba1, and DAPI staining (Sigma, #D9542-10MG). Sections were mounted onto charged glass slides (Genesee Scientific, #30-185) with mounting medium (Invitrogen, 00-4958-02). Images were acquired using a fluorescence microscope (Nikon ECLIPSE Ti2. Objective: 4x; excitation wavelengths: 470 nm, 555 nm, 640 nm and 395 nm; emission wavelengths: 530.5 nm, 615.0 nm, 701.5 nm and 454.0 nm for Alexa Fluor 488, Alexa Fluor 568, Alexa Fluor 647 and DAPI, respectively).

To analyze the co-immunostaining of C05-05 and pS129, mice injected with PFFs in M1 at 3 MPI were injected with C05-05, euthanized at 90 minutes, and brains were flash frozen, sectioned, and stained with S129-phosphorylated α-syn (as above). Z-stacks of ROIs from brain sections were obtained using 2P microscopy. We estimated and extracted the background of 2P maximal fluorescence projection images in MATLAB using the function “imopen” by morphological opening with a disk-shaped structuring element with a manually-adjusted radius in pixels for each image. We then smoothed the resulting image using the function “imgaussfilt” (σ = 1 pixel) and binarized it before analysis. We then performed an object-based colocalization analysis using binarized images. To merge fragmented aggregates and nearby signals, binary masks were subjected to a morphological closing operation using a disk-shaped structuring element with manually-adjusted radius in pixels for each image. We subsequently identified connected components by using the function “bwconncomp”. Colocalization was evaluated in both directions. An object was considered colocalized if at least one pixel in C05-05 labeled image overlapped with an object in the pS129 labeled image. The proportion of colocalized objects was calculated as the number of overlapping objects divided by the total number of objects in each channel. For each overlapping object, the overlap fraction was calculated as the ratio of overlapping pixels to the total area of the object. The percentage of overlap fractions were then computed across all colocalized objects.

### Quantification of S129-phosphorylated **α**-syn pathology

Immunofluorescence images were registered to the Allen Brain Atlas CCFv3 using the QUINT workflow^56^ to calculate the density of S129-phosphorylated α-syn across brain regions. An RBG image of each section was exported from ImageJ as an 8-bit PNG and JPEG. This conversion involved down sampling by a factor of approximately 5.7. The JPEG images, grouped by mouse identification, were used for spatial registration in QuickNII. Brain images were uploaded in Filebuilder and saved as an XML file to be compatible with QuickNII that could then be used to register each brain slice image to its appropriate anatomical location and orientation within the Allen Brain Atlas CCFv3. After this initial spatial registration further adjustments for a precision fit to the atlas were made in VisuAlign. Object segmentation was performed using Ilastik. We trained and validated a custom Ilastik model and used it to segment α-syn pathology. Object segmentations and their atlas registration files were then processed using Nutil to assign objects to brain regions. Final quantification of the density of α-syn pathology as a proportion of the total area of each brain region of interest was then calculated using customized MATLAB scripts.

### Statistical Analyses

All data are plotted as mean +/− standard error of the mean (s.e.m), unless otherwise stated. Sample sizes were not based on *a priori* power calculations but are consistent with, or greater than, other studies in the field using similar techniques, including our own^51,57^. Statistical analyses were performed in MATLAB. Data were analyzed using parametric or non-parametric statistical tests, as indicated in the figure legends. To account for nested data (measuring multiple cells nested within individual mice) we used mixed effects models^58^. We calculated intraclass correlation coefficients (ICCs) for OSI, DSI, peak amplitude, calcium transient frequency, and AUC and found that across these measures, ICC values were generally low (<0.5) (**Tables. S1-5**), consistent with low correlation due to clustering of neurons within individual mice and thus large effective sample sizes. For experiments measuring percentage of visually responsive cells or percentage of cells with OSI/DSI > 0.6 (**Fig. 2f,i,j**), we used GLME models with a binomial distribution, with individual mouse ID modeled as a random effect. For other longitudinal analyses we used linear mixed effects (LME) models, with cells nested within individual mice as a random effect. For models with more than one fixed effect, ANOVA was used to test for overall significance of fixed effects. For fixed effects that achieved significance at the level of the ANOVA, *p* values for individual coefficients from the model were corrected for multiple hypothesis testing using the Benjamini & Hochberg procedure^59^ (# and *, *p* < 0.05; ## and **, *p* < 0.01; ### and ***, *p* < 0.001). In the figures, results are presented with *p* values for overall fixed effects shown as an inset on respective graphs, with *p* values for individual coefficients depicted adjacent to the respective data points where appropriate. The analysis code used in this study is publicly available at our laboratory’s GitHub repository (https://github.com/zeigerlab/PD-aSyn-Visual-Processing). The results of all statistical tests are provided in the Supplementary File.

## Declarations

## Author Contributions

ATT designed and performed research, analyzed data, and wrote the manuscript; NC performed research and analyzed data; SZ performed research; NZ performed research; RM performed research; KI performed research and analyzed data; MH designed research, analyzed data, and provided C05-05; CP designed research; WZ designed and performed research, analyzed data, and wrote the manuscript. All authors have approved the final version of the manuscript and consent to its submission.

## Ethics approval and consent to participate

All experiments followed the US National Institutes of Health guidelines for animal research, under an animal use protocol approved by the University of California, Los Angeles Animal Research Committee. Other ethical approval was waived due to the nature of the study or the retrospective nature of the study.

### Conflict of Interest

The authors declare no competing financial or non-financial interests.

## Acknowledgements

We thank Ms. Brenda Vasquez, Ms. Callie Liu, and Ms. Sravya Gadepalli for assistance with mouse breeding and colony management. We thank Ms. Britney Burnasky and Ms. Ashley Cao for assistance with mouse behavior testing. We thank Drs. Baruc Campos, Sammy Alhassen, and Carlos Portera-Cailliau for helpful feedback, constructive criticism, and advice.

This work was supported by National Institutes of Health grant 1R01NS129517-01, American Parkinson’s Disease Association research grant 1178689 and Dr. George C. Cotzias Memorial Fellowship 1268719, the UCLA Laurie and Steven C. Gordon Parkinson’s Disease Grant Program, and the UCLA David Binder Fund for Lewy Body Disease Research.

## Data Availability

Statistical results in the study are included in Supplementary Data File. The analysis codes and data used in this study are publicly available at our laboratory’s GitHub repository (https://github.com/zeigerlab/PD-aSyn-Visual-Processing).

## Supplementary Figures

**Supplementary Figure 1:**
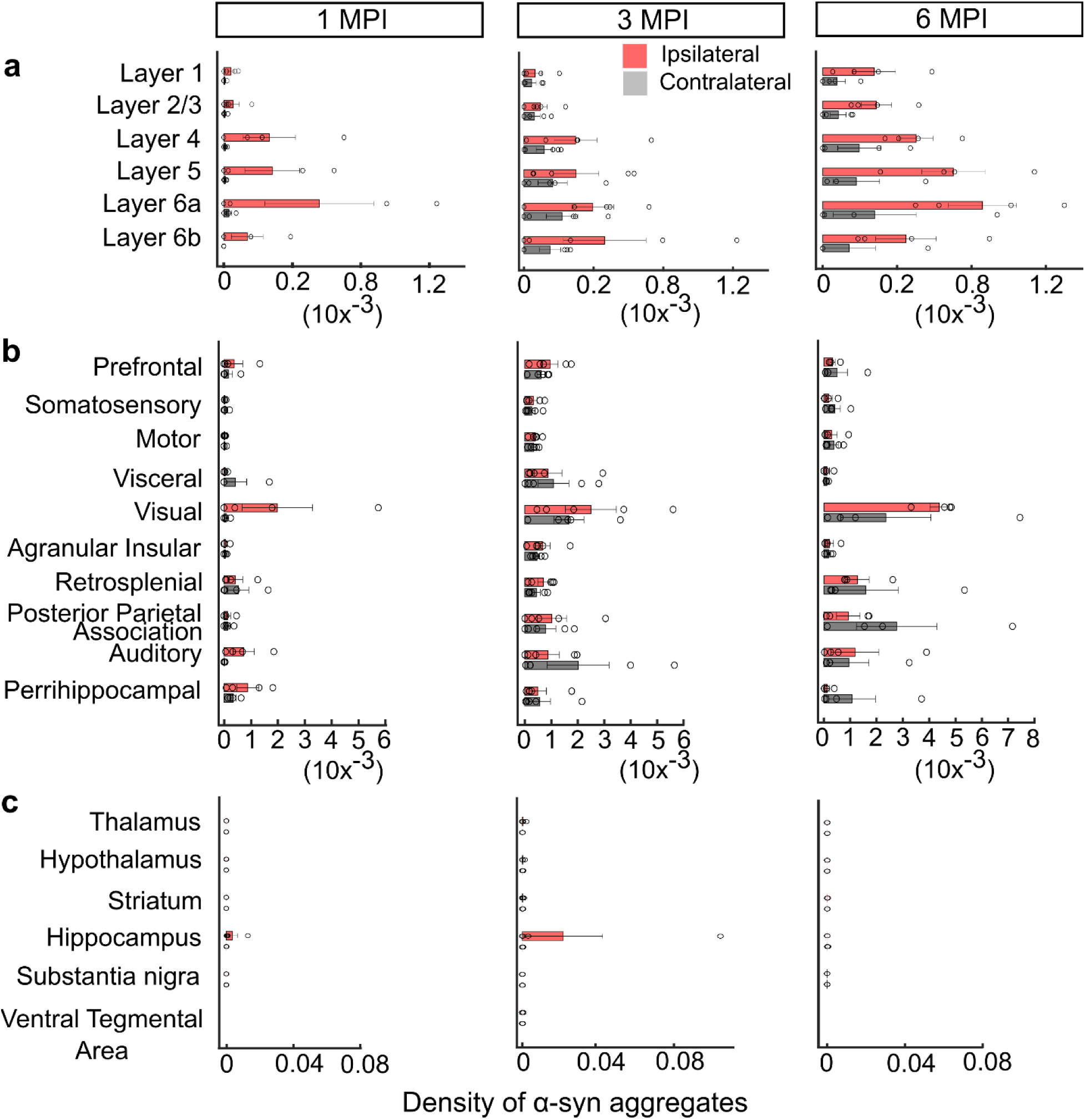
Distribution of pS129^+^ α-syn aggregates in V1 and other brain regions over time. **a-c :** Density of pS129^+^ α-syn aggregates in each layer of ipsilateral (red) and contralateral (gray) visual cortical areas *(a)*, other cortical regions *(b)* and other brain regions *(c).* Bars show mean ± s.e.m., with data points for individual mice (n=5 mice) overlaid.

**Supplementary Figure 2:**
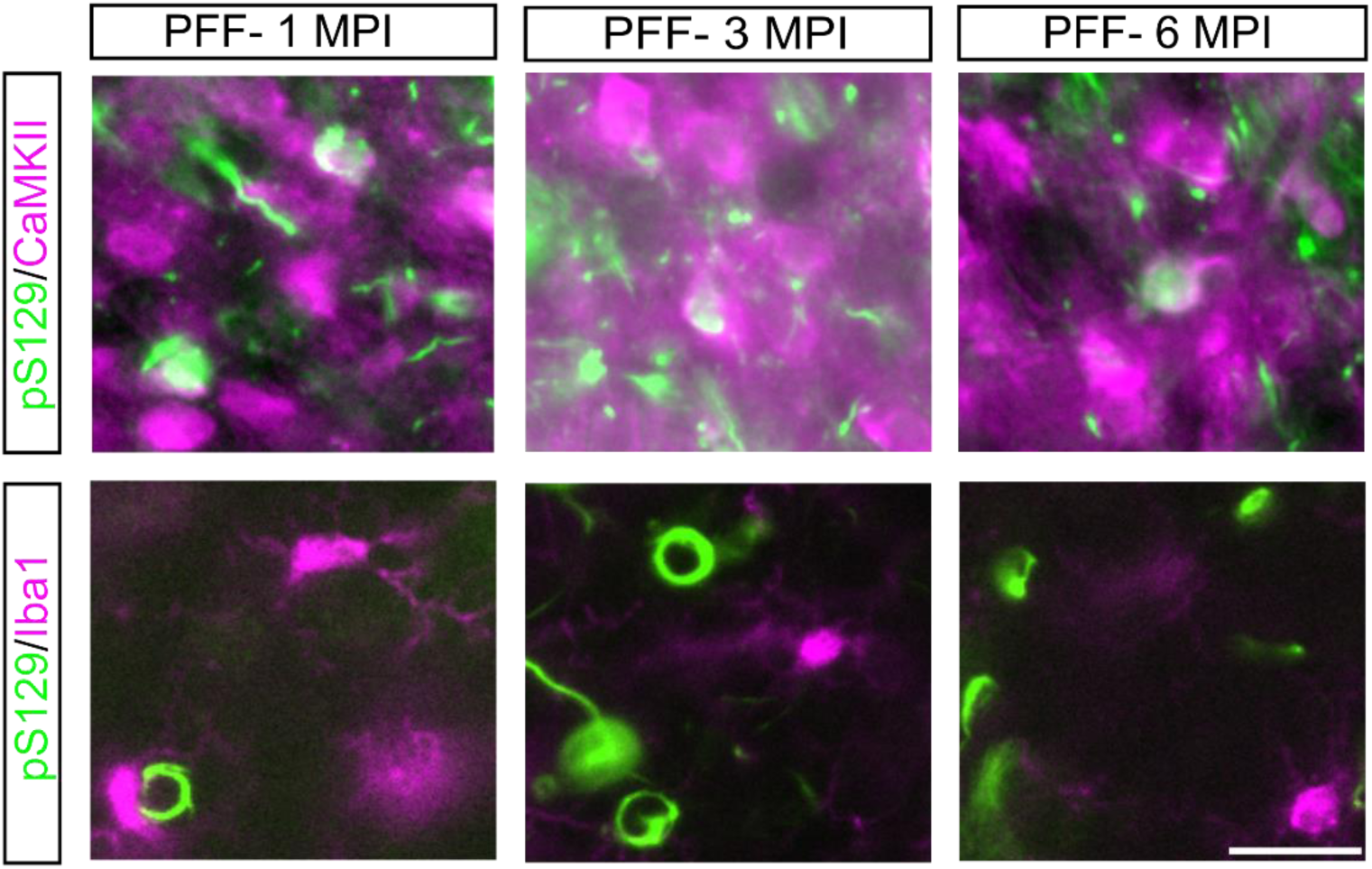
pS129^+^ α-syn aggregates predominately colocalize with excitatory neurons in V1. Representative immunofluorescence composite images showing co-staining of pS129^+^ aggregates with CaMKII^+^ (Top Row) or Iba1^+^ (Bottom Row) cells in PFF-injected mice in V1 at 1, 3, and 6 MPI. pS129^+^ α-syn aggregates were frequently observed within CaMKII^+^ excitatory neurons across all time points, whereas little to no colocalization of pS129^+^ aggregates with Iba1^+^ microglia was detected. Scale bar, 20 μm.

**Supplementary Figure 3:**
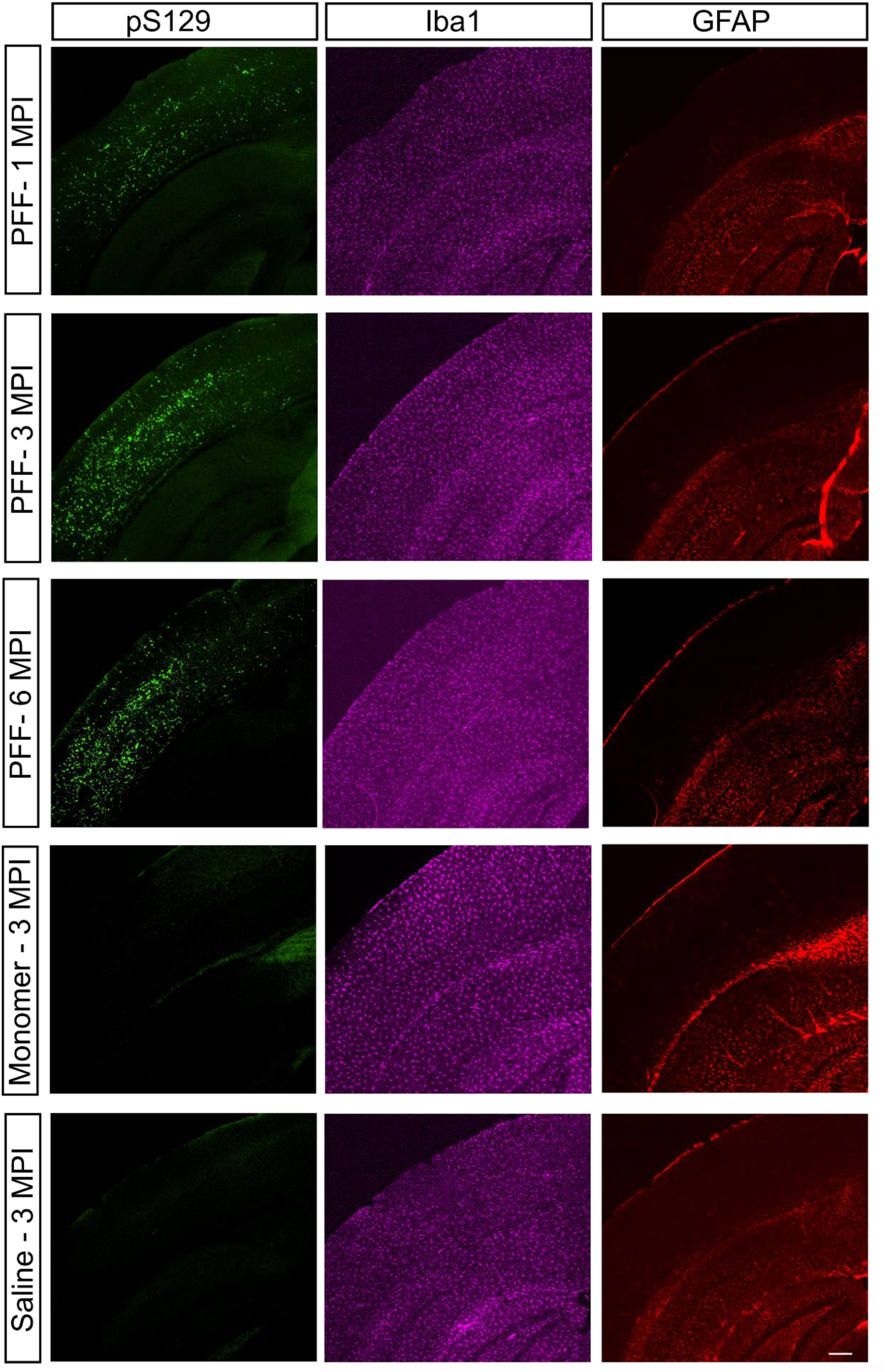
Qualitatively similar GFAP and Iba1 staining in V1 after injection of PFFs, α-syn monomers, or saline. Representative immunofluorescence composite images of pS129^+^ aggregates, Iba1^+^ microglia, and GFAP^+^ astrocytes in mice injected with α-syn PFFs, α-syn monomers, or saline in V1. Robust pS129^+^ α-syn aggregates were observed in PFF-injected mice, whereas α-syn monomer-injected and control mice showed no detectable pS129^+^ aggregates. Iba1^+^ and GFAP^+^ cells exhibited comparable density and expression across groups, with no overt qualitative differences in α-syn PFF- or monomer-injected mice relative to saline controls. Scale bar, 20 μm.

**Supplementary Figure 4:**
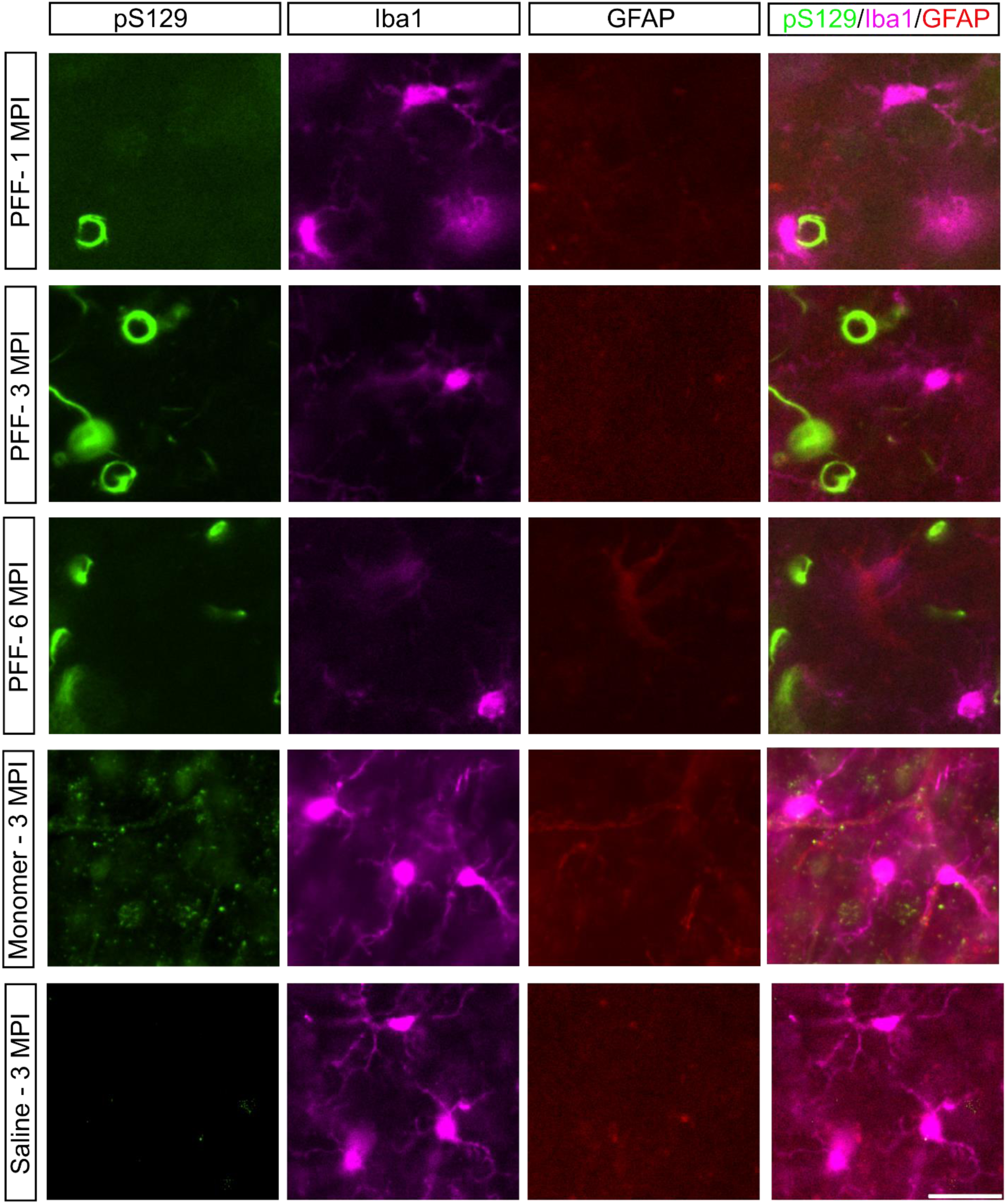
Injection of PFFs, α-syn monomers, and saline results in comparable glial responses in V1. Representative immunofluorescence composite images of pS129^+^ aggregates, Iba1^+^ microglia, and GFAP^+^ astrocytes in mice injected with α-synuclein PFFs, α-syn monomers, or saline in V1. Robust pS129^+^ α-syn aggregates were observed in PFF-injected mice, but not α-syn monomer-injected or saline-injected control mice. Iba1^+^ microglia exhibited comparable distribution and morphology across groups. There were few GFAP^+^ astrocytes in V1, with no overt evidence of gliosis in α-syn PFF- or monomer-injected mice relative to saline controls. Scale bar, 20 μm.

**Supplementary Figure 5:**
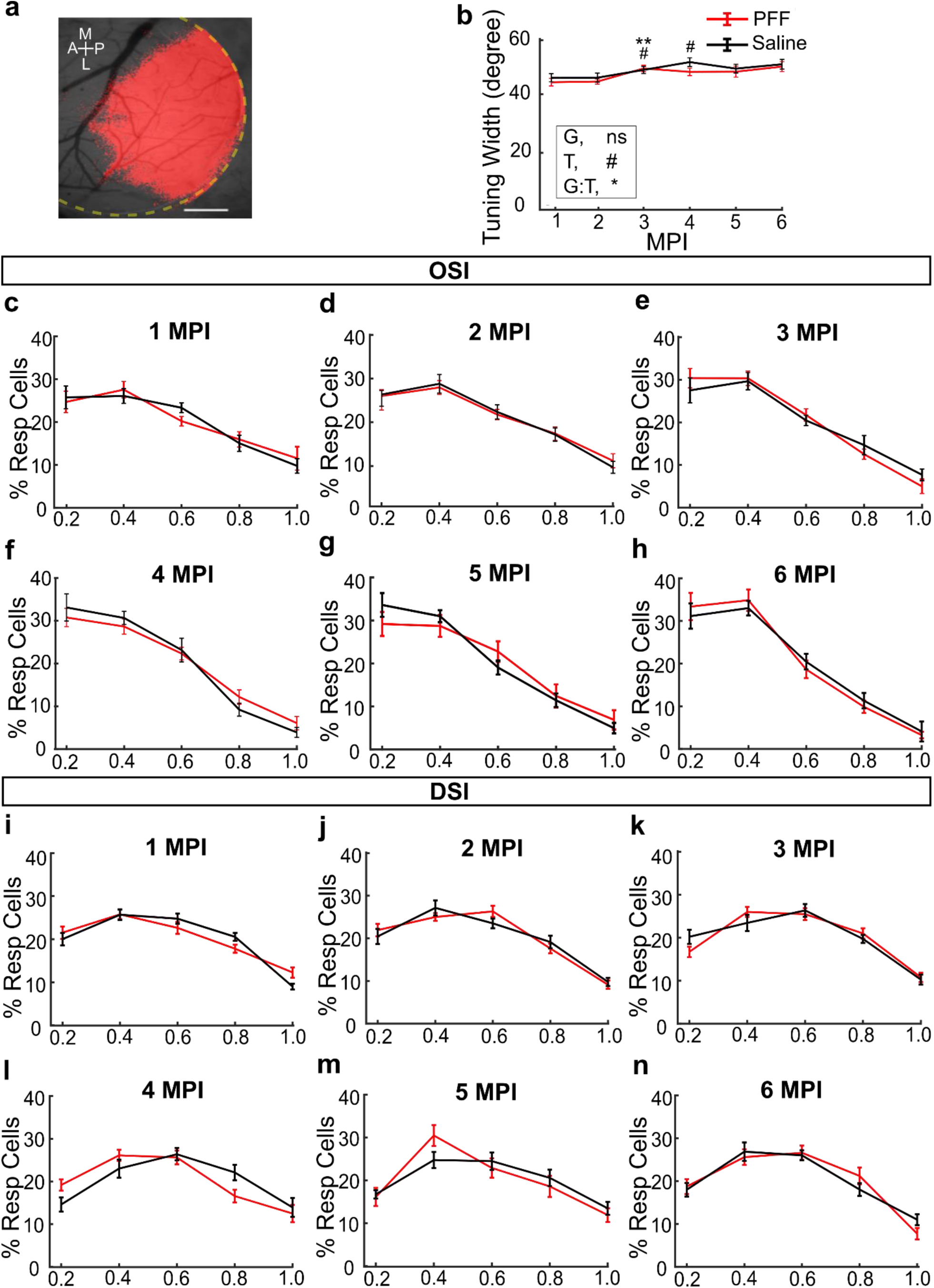
Changes in proportions of orientation and direction selective cells over time. **a:** Representative IOSI map of visually evoked neuronal activity overlaid on an image of the cortical vasculature through the cranial window. The dotted yellow line marks the edge of the cranial window. Scale bar. 0.5 mm. **b:** Tuning width of L2/3 neurons in V1 for control and PFF-injected mice over time. LME model, ANOVA for fixed effects of group (G, *p* = 0.124), timepoint (T, #, *p* = 0.0126), and group-by-timepoint interaction (G:T, *, *p* = 0.0240). Significance for individual coefficients for timepoint (T) or group-by-timepoint interaction (G:T), corrected using Benjamini and Hochberg’s (B & H) method, are indicated over corresponding data points (# and *, *p* < 0.05; ## and **, *p* < 0.01; ### and ***, *p* < 0.001). **c-h:** Distributions of the OSIs for all visually responsive neurons recorded in control and PFF-injected mice at 1-6 MPI. **i-n:** Distributions of the DSIs for all visually responsive neurons recorded in control and PFF-injected mice at 1-6 MPI.

**Supplementary Figure 6:**
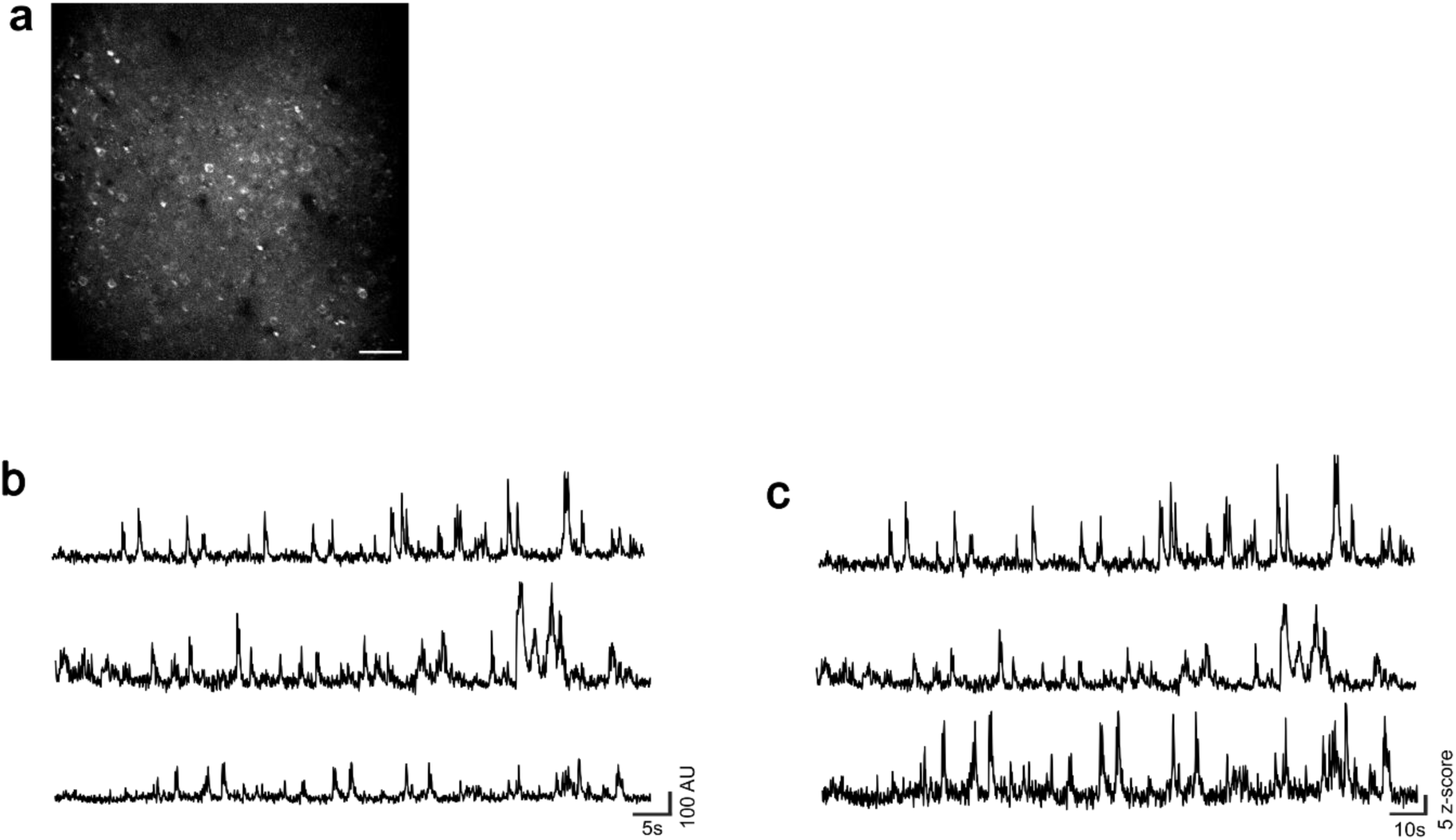
Stable jRGECO1a expression and representative calcium activity traces in control mice. **a:** Representative maximum intensity projection image of L2/3 cortical neurons expressing jRGECO1a at 6 MPI, demonstrating absence of filled cells typical of calcium indicator toxicity. Scale bar, 50 μm. **b-c:** Representative raw (*b*) and z-scored (*c*) jRGECO1a fluorescence traces from 3 individual neurons recorded in control mice at 3 MPI.

**Supplementary Figure 7:**
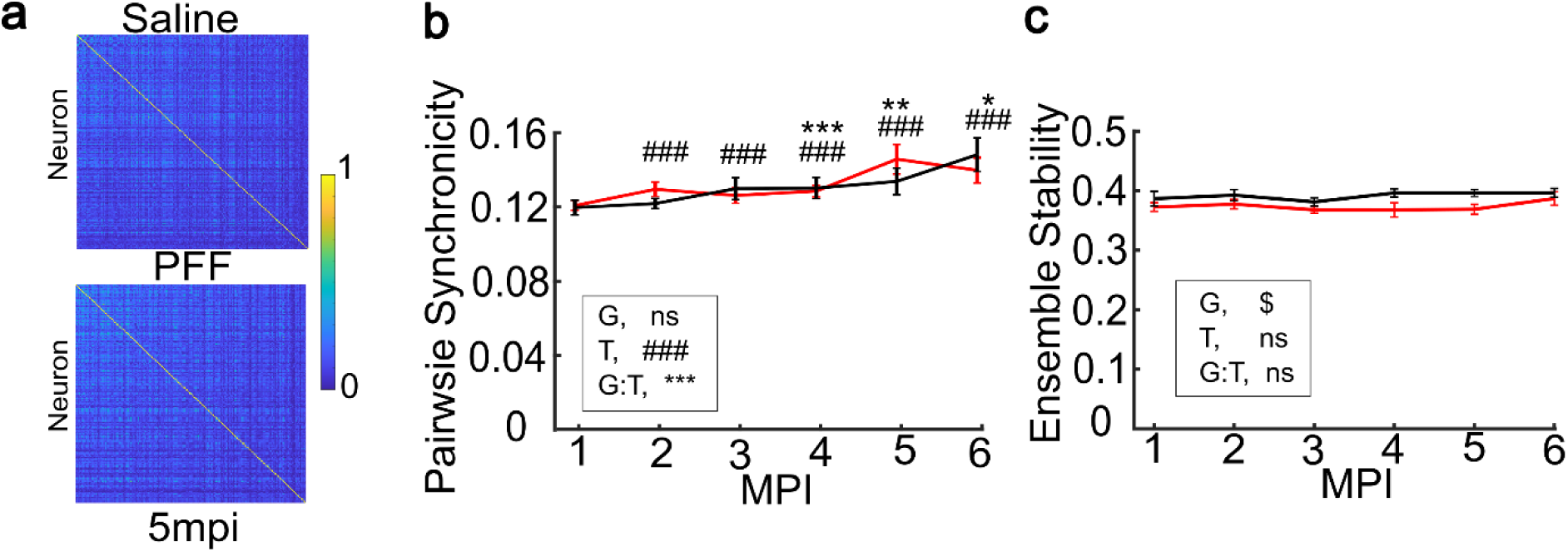
α-syn pathology causes minimal change in neuronal synchronicity and ensemble stability of visually evoked activity. **a**: Correlograms of pairwise correlations of visually evoked calcium transients from PFF-injected or control mice at 5 MPI. The scale denotes the SØrenson-Dice correlation coefficient. **b:** Pairwise synchronicity of neurons during visually evoked activity in control and PFF-injected mice. LME model, ANOVA for fixed effects of group (G, *p* = 0.239), timepoint (T, ###, *p* = 1.50 × 10^−141^), and group-by-timepoint interaction (G:T, ***, *p* = 6.60 × 10^−12^). Significance for individual coefficients for timepoint (T) or group-by-timepoint interaction (G:T), corrected using B & H method, are indicated over corresponding data points in *b-c* ($, # and *, *p* < 0.05; ## and **, *p* < 0.01; ### and ***, *p* < 0.001). Individual timepoints show means ± s.e.m for data in *b-c.* **c:** Stability of neuronal ensembles during visually evoked activity in control and PFF-injected mice. LME model, ANOVA for fixed effects of group (G, $, *p* = 0.00214), timepoint (T, *p* = 0.0642), and group-by-timepoint interaction (G:T, *p* = 0.121).

**Supplementary Figure 8:**
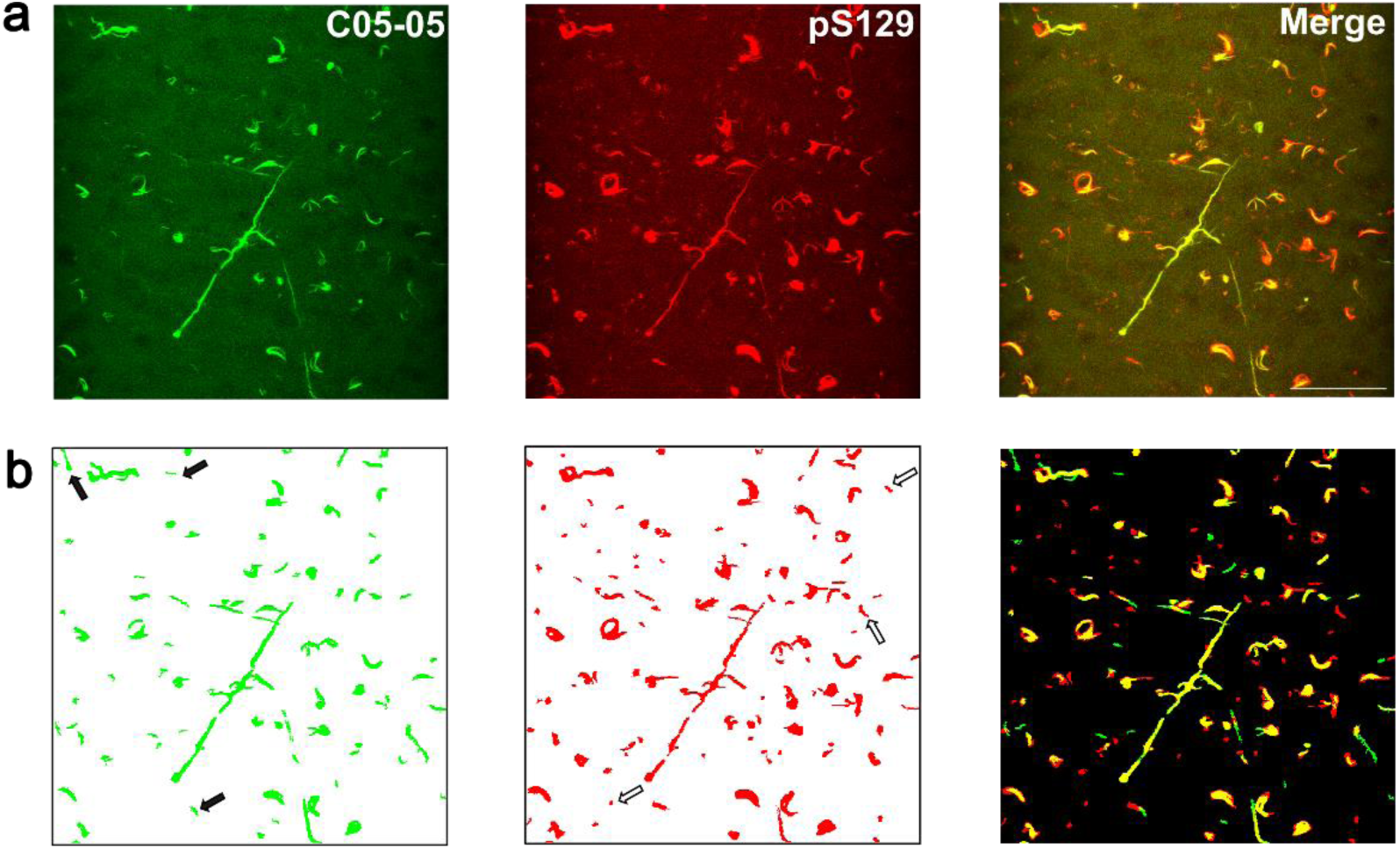
C05-05 selectively binds S129-phosphorylated α-synuclein aggregates. **a:** Representative 2P images of C05-05 (left), pS129 (middle) and the raw composite image (right) in ex vivo coronal tissue sections from a PFF-injected mouse. Scale bar = 50 μm. **b**: Corresponding binarized images of C05-05 (left), pS129 (middle) and the binarized composite image (right). Solid black arrows denote false positives (C05-05^+^, pS129^-^), while hollow black arrows highlight false negatives (C05-05^-^, pS129^+^).

**Supplementary Figure 9:**
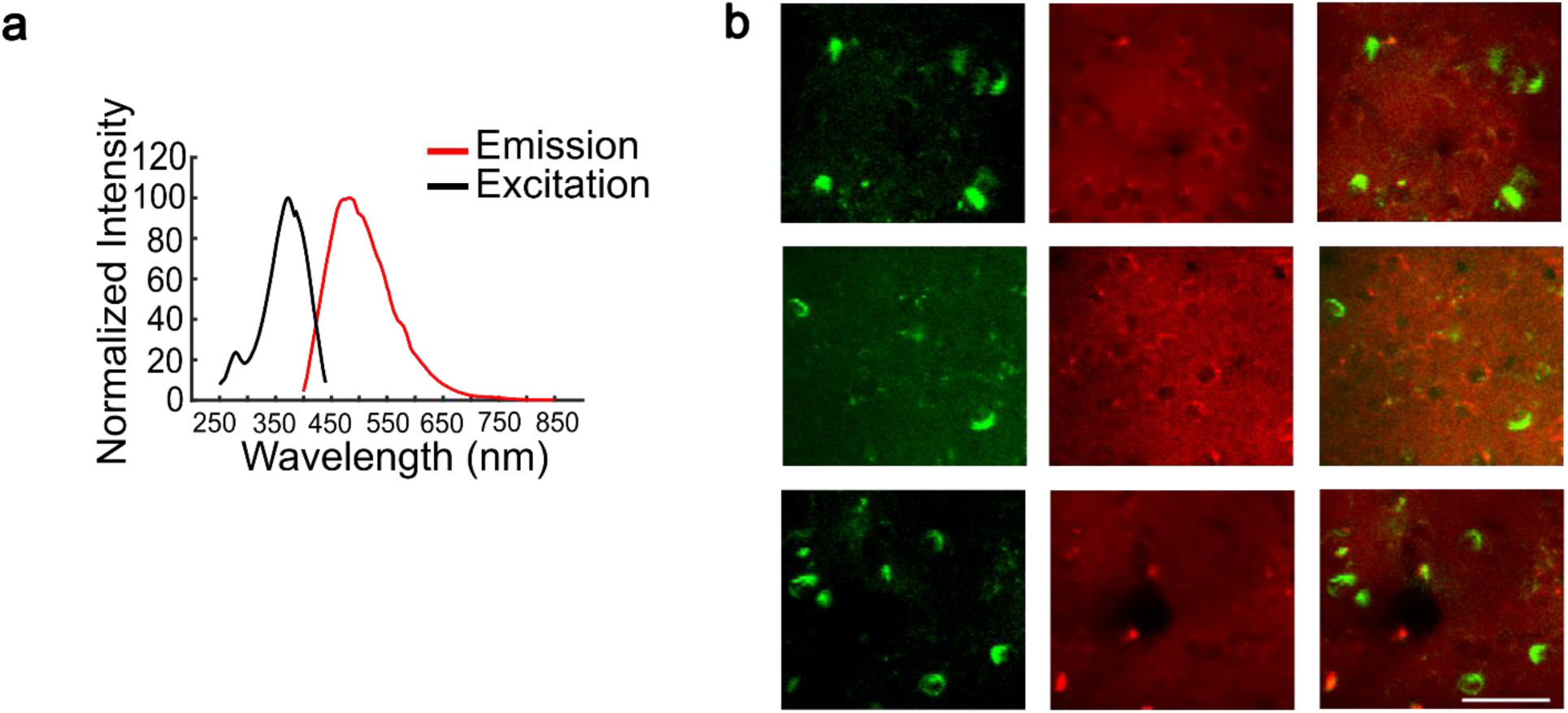
C05-05 and jRGECO1a fluorescence signals are separable. **a:** Excitation and emission spectrum of C05-05. **b:** Representative in vivo 2P images of C05-05 (left), jRGECO1a (middle) and the composite image (right) in PFF-injected mice at 6 MPI. Scale bar = 50 μm.

**Supplementary Figure 10:**
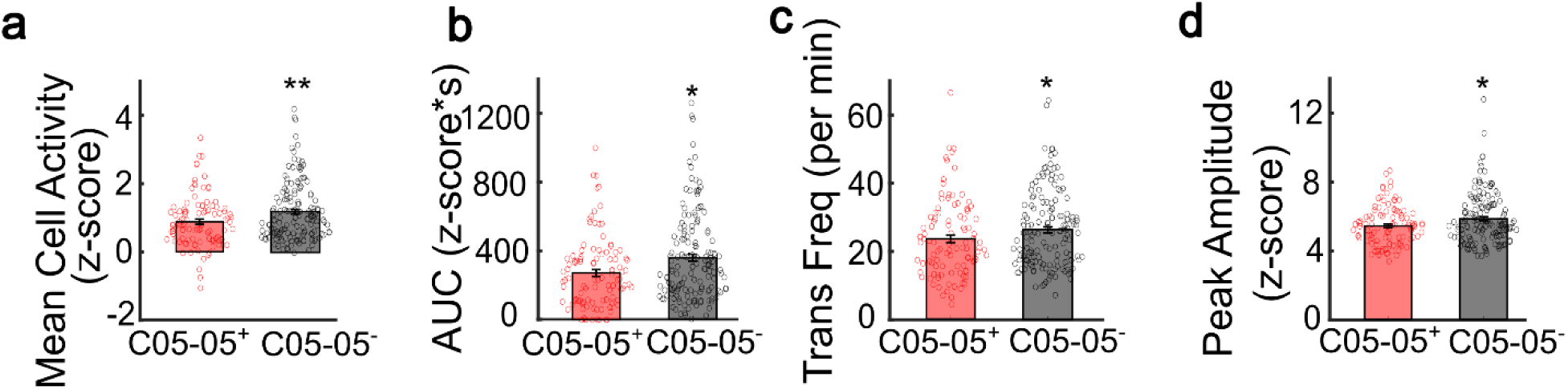
Intracellular α-syn inclusions are associated with reduced spontaneous activity in neurons in V1 of PFF-injected mice. **a-d :** Mean z-scored fluorescence (*a*), AUC (*b*), transient frequency (*c*), and transient peak amplitude (*d*) in C05-05^+^ (red, n = 100 cells) and C05-05^-^ (gray, n = 153 cells) neurons from PFF-injected mice under spontaneous conditions at 4 and 5 MPI. Two-sample, two-tailed *t* test, *p* = 0.0036 (*a*), *p* = 0.0056 (*b*), *p* = 0.041 (*c*) and *p* = 0.015 (*d*). Bars show mean ± s.e.m., with data points for individual neurons overlaid.

**Supplementary Figure 11:**
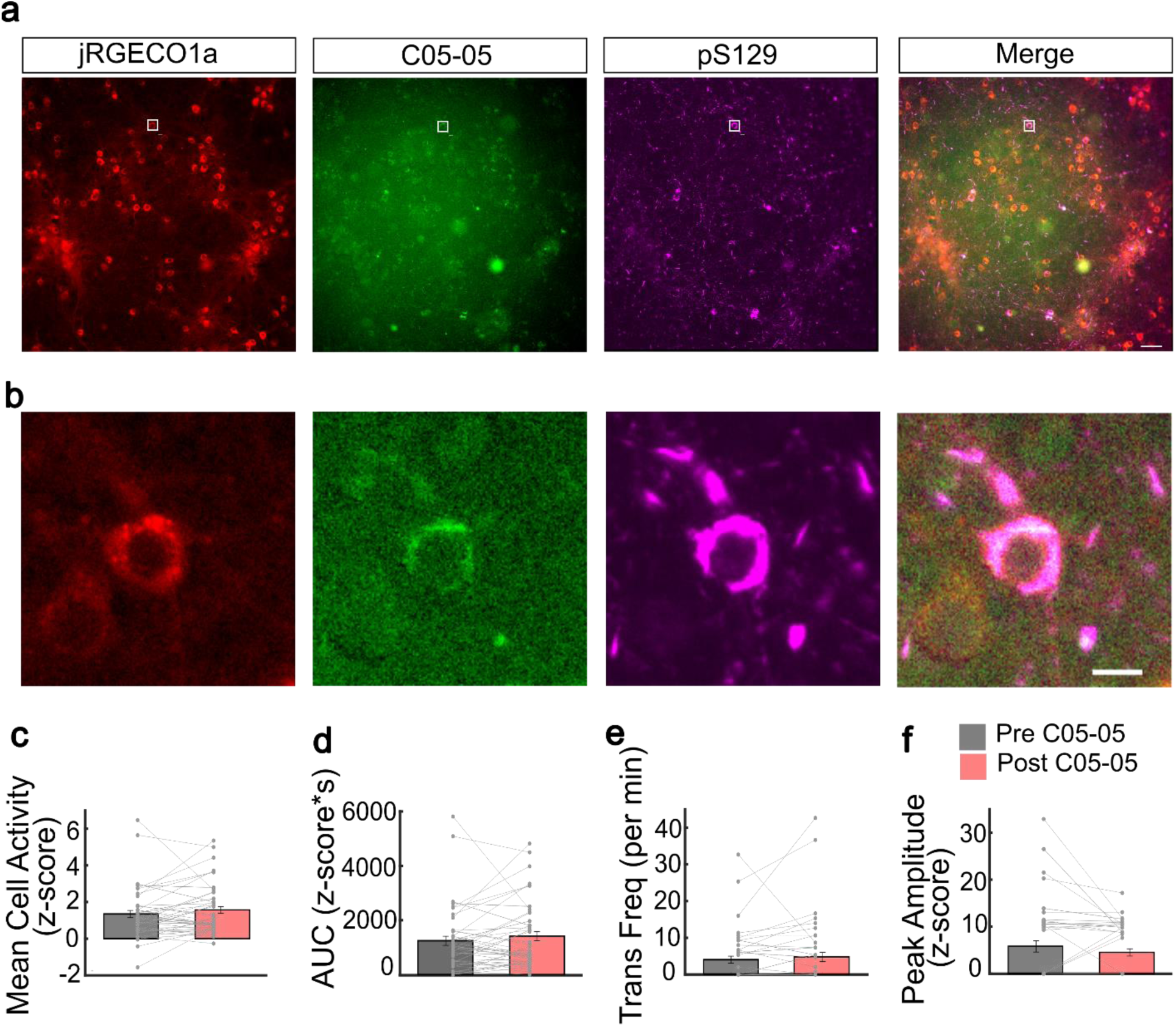
C05-05 does not significantly alter neuronal activity measured by jRGECO1a in PFF-seeded primary cortical neurons. **a:** A representative FOV showing jRGECO1a (left), C05-05 (second from left), pS129 (third from left) and the composite image (right) in a PFF-seeded well. Boxes indicate a representative neuron which is jRGECO1a^+^, C05-05^+^ and pS129^+^. Scale bar = 50 μm. **b:** Higher magnification images of the neuron from (*a)* expressing jRGECO1a (left), C05-05 (middle) and its composite image (right). Scale bar = 10 μm. **c-j:** Mean z-scored fluorescence *(c)*, AUC *(d)*, transient frequency *(e)*, and transient peak amplitude *(f)* of jRGECO1a^+^/C05-05^+^/ pS129^+^ primary cortical neurons before (Pre C05-05) and after C05-05 treatment (Post C05-05). Paired Student’s *t* test, *p* = 0.157 *(c)*, *p* = 0.214 *(d)*, *p* = 0.107 *(e)*, *p* = 0.015 *(f)*. Bars show mean ± s.e.m., with data points for individual neurons and connecting lines for paired measurements from the same neuron overlaid.

**Supplementary Figure 12:**
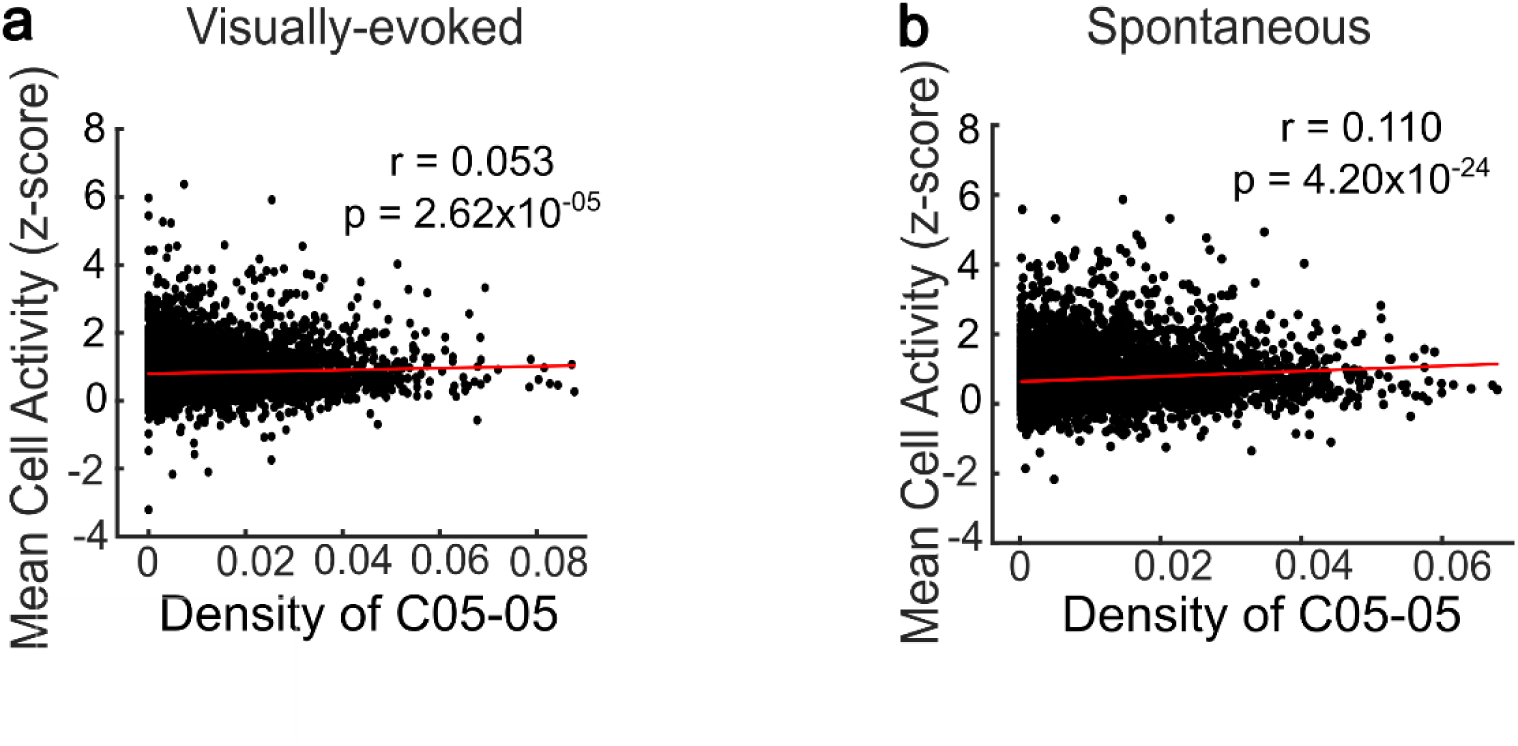
Local burden of α-syn pathology is associated with increased activity of inclusion-free neurons in V1. **a-b:** Correlation between mean z-score of calcium fluorescence traces of individual C05-05^-^ neurons and local C05-05 burden within a radius of 100 pixels (83 µm) from PFF-injected mice under visually evoked (*a*) and spontaneous (*b*) conditions at 4-5 MPI. Pearson correlation coefficient (r = 0.053, *p* = 2.62 × 10^−5^, n = 6264 neurons in *a* and r = 0.110, *p* = 4.20 × 10^−24^, n = 8470 neurons in *b*).

**Supplementary Figure 13:**
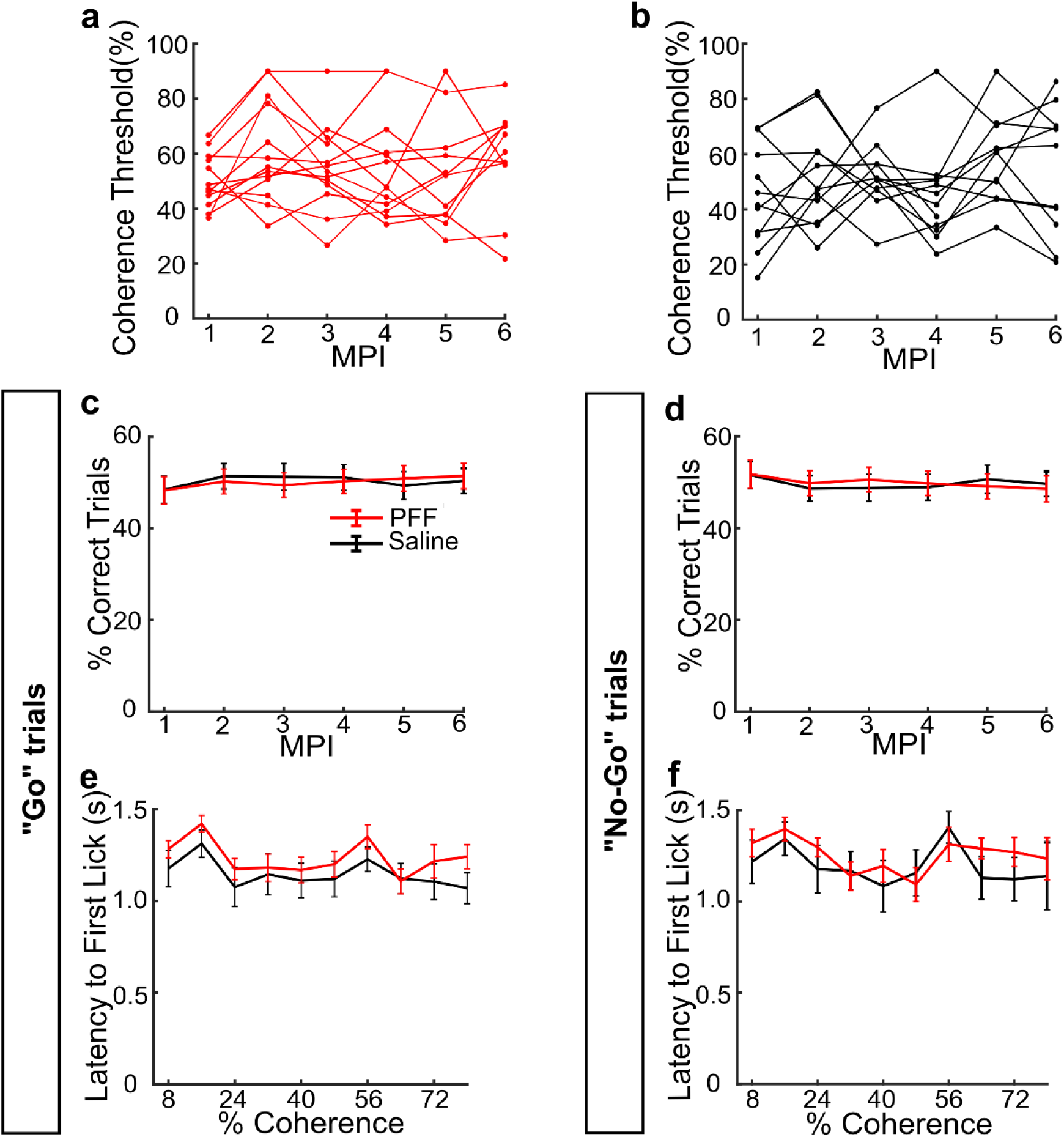
Changes in performance of individual mice and behavioral performance metrics in the motion discrimination task. **a-b:** Coherence thresholds over time up to 6 MPI in each PFF-injected *(a)* and control *(b)* mouse. **c-d:** Performance across “Go” *(c)* and “No-Go” *(d)* trials over time up to 6 MPI in PFF-injected and control mice. **e-f:** Average latency to first lick after stimulus presentation across test coherence values at 4 and 5 MPI in “Go” *(e)* and “No-Go” *(f)* trials in PFF-injected and control mice.

**Supplementary Figure 14:**
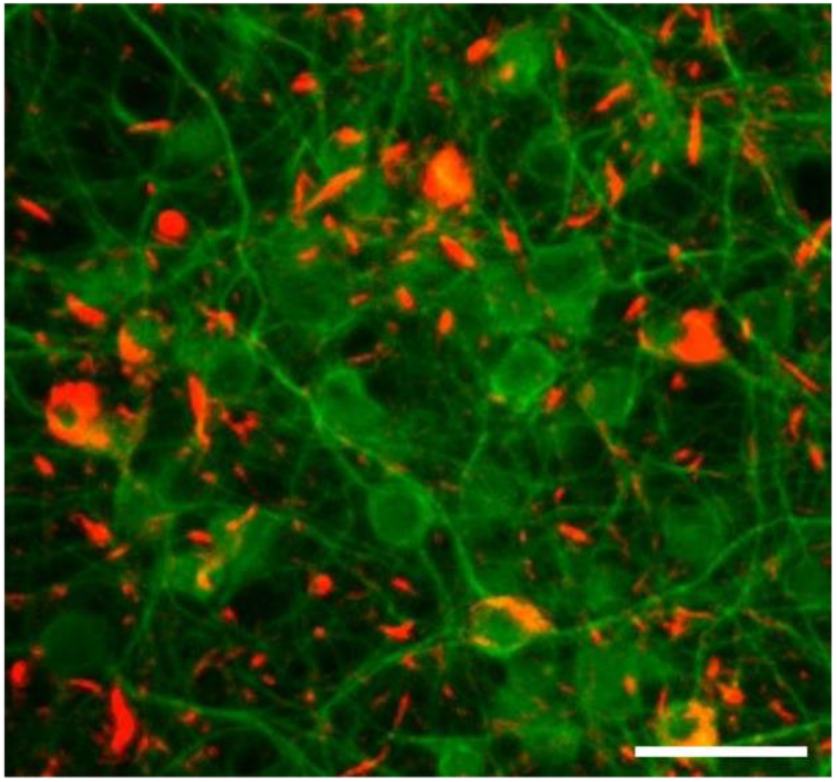
Validation of seeding efficacy of PFFs. Composite immunostaining image of pS129 (Red) and MAP2 (Green) in a PFF-seeded primary neuron culture. Scale bar = 20 μm.

## Supplementary Tables

**Supplementary Table 1:** ICC of OSI from PFF-injected or control mice (Mean = 0.10)

| Group | MPI | # mice | #cells | ICC |
| --- | --- | --- | --- | --- |
| Saline | 1 | 19 | 2062 | 0.11 |
| Saline | 2 | 19 | 2503 | 0.07 |
| Saline | 3 | 17 | 1573 | 0.12 |
| Saline | 4 | 14 | 1352 | 0.08 |
| Saline | 5 | 16 | 1171 | 0.06 |
| Saline | 6 | 12 | 978 | 0.08 |
| PFF | 1 | 22 | 2153 | 0.16 |
| PFF | 2 | 24 | 2568 | 0.12 |
| PFF | 3 | 23 | 2233 | 0.07 |
| PFF | 4 | 15 | 1507 | 0.05 |
| PFF | 5 | 16 | 1022 | 0.13 |
| PFF | 6 | 12 | 1045 | 0.13 |

**Supplementary Table 2:**
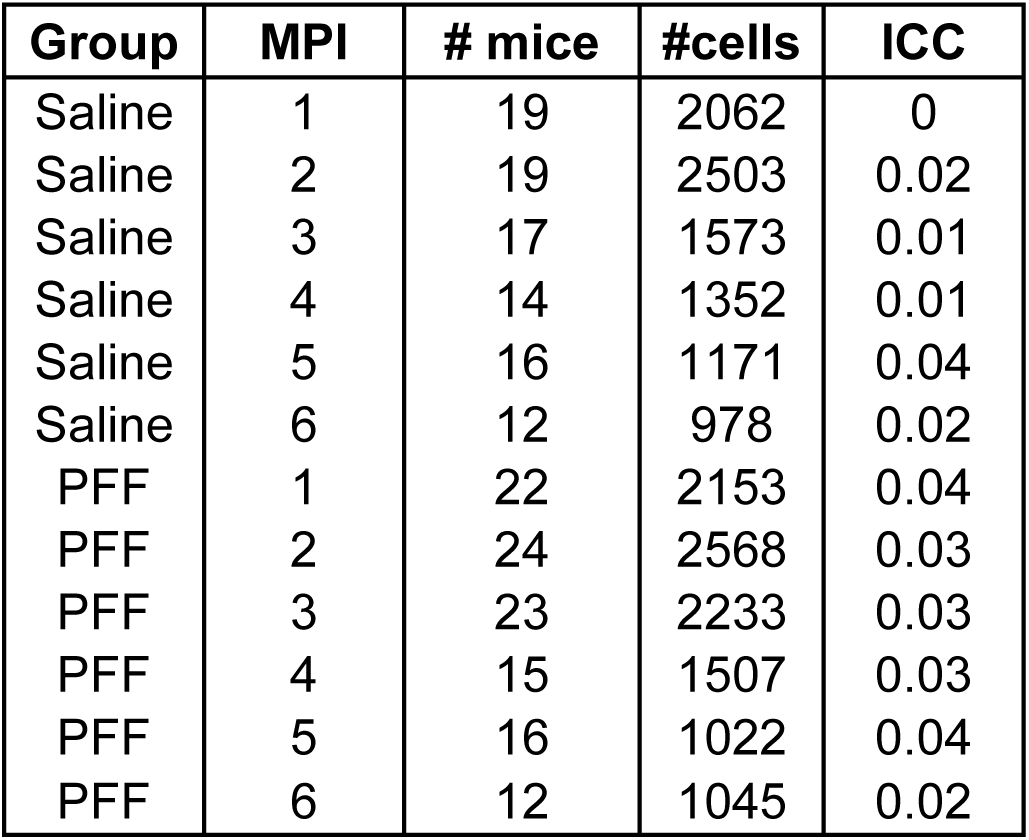
ICC of DSI from PFF-injected or control mice (Mean = 0.02)

**Supplementary Table 3:**
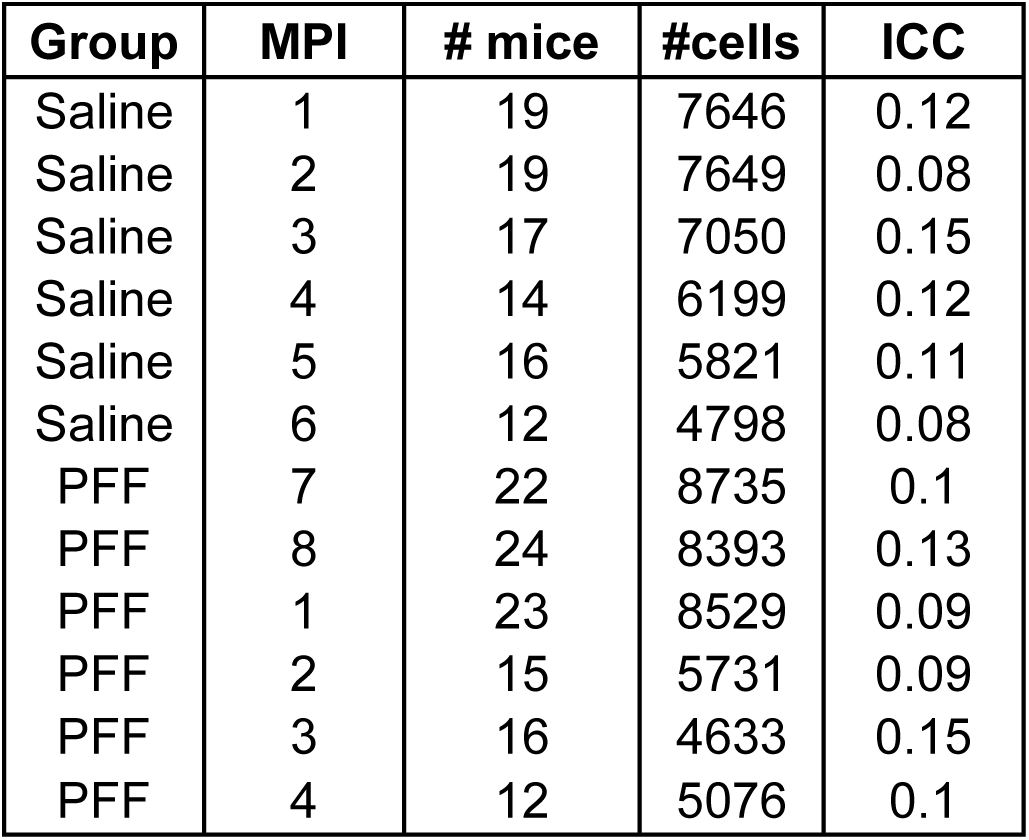
ICC of peak amplitude of calcium transients from PFF-injected or control mice (Mean = 0.10)

| Group | MPI | # mice | #cells | ICC |
| --- | --- | --- | --- | --- |
| Saline | 1 | 19 | 7646 | 0.12 |
| Saline | 2 | 19 | 7649 | 0.08 |
| Saline | 3 | 17 | 7050 | 0.15 |
| Saline | 4 | 14 | 6199 | 0.12 |
| Saline | 5 | 16 | 5821 | 0.11 |
| Saline | 6 | 12 | 4798 | 0.08 |
| PFF | 7 | 22 | 8735 | 0.1 |
| PFF | 8 | 24 | 8393 | 0.13 |
| PFF | 1 | 23 | 8529 | 0.09 |
| PFF | 2 | 15 | 5731 | 0.09 |
| PFF | 3 | 16 | 4633 | 0.15 |
| PFF | 4 | 12 | 5076 | 0.1 |

**Supplementary Table 4:** ICC of frequency of calcium transients from PFF-injected or control mice (Mean = 0.44)

| Group | MPI | # mice | #cells | ICC |
| --- | --- | --- | --- | --- |
| Saline | 1 | 19 | 7646 | 0.5 |
| Saline | 2 | 19 | 7649 | 0.51 |
| Saline | 3 | 17 | 7050 | 0.52 |
| Saline | 4 | 14 | 6199 | 0.5 |
| Saline | 5 | 16 | 5821 | 0.49 |
| Saline | 6 | 12 | 4798 | 0.45 |
| PFF | 7 | 22 | 8735 | 0.52 |
| PFF | 8 | 24 | 8393 | 0.49 |
| PFF | 1 | 23 | 8529 | 0.52 |
| PFF | 2 | 15 | 5731 | 0.47 |
| PFF | 3 | 16 | 4633 | 0.47 |
| PFF | 4 | 12 | 5076 | 0.56 |

**Supplementary Table 5:** ICC of AUC from the entire spontaneous activity trace from neurons from PFF-injected or control mice (Mean = 0.03)

| Group | MPI | # mice | #cells | ICC |
| --- | --- | --- | --- | --- |
| Saline | 1 | 19 | 7646 | 0.02 |
| Saline | 2 | 19 | 7649 | 0.01 |
| Saline | 3 | 17 | 7050 | 0.03 |
| Saline | 4 | 14 | 6199 | 0.02 |
| Saline | 5 | 16 | 5821 | 0.03 |
| Saline | 6 | 12 | 4798 | 0.05 |
| PFF | 7 | 22 | 8735 | 0.01 |
| PFF | 8 | 24 | 8393 | 0.04 |
| PFF | 1 | 23 | 8529 | 0.03 |
| PFF | 2 | 15 | 5731 | 0.07 |
| PFF | 3 | 16 | 4633 | 0.04 |
| PFF | 4 | 12 | 5076 | 0.05 |

## Notes

### Competing Interest Statement

The authors have declared no competing interest.

### Summary of Updates

We have updated figure 1 and added multiple supplementary figures.

